# Ribosome heterogeneity in *Drosophila melanogaster* gonads through paralog-switching

**DOI:** 10.1101/2020.01.20.913020

**Authors:** Tayah Hopes, Karl Norris, Michaela Agapiou, Charley G.P. McCarthy, Philip A. Lewis, Mary J O’Connell, Juan Fontana, Julie L Aspden

**Author notes:** These authors contributed equally to this paper (Tayah Hopes, Karl Norris).

## Abstract

Ribosomes have long been thought of as homogeneous macromolecular machines, but recent evidence suggests they are heterogeneous and could be specialised to regulate translation. Here, we have characterised ribosomal protein heterogeneity across 4 tissues of *Drosophila melanogaster*. We find that testes and ovaries contain the most heterogeneous ribosome populations, which occurs through a combination of paralog-enrichment and paralog-switching. We have solved structures of ribosomes purified from *in vivo* tissues by cryo-EM, revealing differences in precise ribosomal arrangement for testis and ovary 80S ribosomes. Differences in the amino acid composition of paralog pairs and their localisation on the ribosome exterior indicate paralog-switching could alter the ribosome surface, enabling different proteins to regulate translation. One testis-specific paralog-switching pair is also found in humans, suggesting this is a conserved site of ribosome heterogeneity. Overall, this work allows us to propose that mRNA translation might be regulated in the gonads through ribosome heterogeneity, providing a potential means of ribosome specialisation.

## INTRODUCTION

Protein synthesis is essential across the tree of life and undertaken by the highly conserved macromolecular complex of “the ribosome”. mRNA translation is regulated at many levels, but until recently the ribosome itself was not thought to be part of this control system. Recent studies have suggested that ribosomes can contribute to gene expression regulation, through specific changes in their composition, i.e. specialisation [1-3]. These “specialised ribosomes” are thought to contribute to the translation of specific mRNA pools; but the mechanism by which this takes place is yet to be understood.

Previous analyses in a variety of organisms (mouse embryonic stem cells [2], yeast [4], and human cell lines [5]) have shown that the composition of ribosomes is heterogeneous. These different ribosome populations may be able to regulate translation. In fact, specialisation of ribosomes is thought to be able to occur through heterogenous ribosomes that contain a) additional protein components [6], b) substitution of ribosomal protein (RP) paralogs [7], c) post-translational modification of RPs [8], and d) rRNA modifications [9]. All these changes to the composition of ribosomes could potentially contribute to functionally specialised ribosomes [10].

Two significant factors have contributed to the logic behind the idea of specialised ribosomes: a) prevalence of tissue specific RP expression and b) distinctive phenotypes when RP genes are disrupted [11]. Many RPs exhibit differences in expression levels across various tissues in mammals [1, 12, 13], plants [14], and insects [7]. For example, RpS5A and RpS5B are expressed in different cell types during early *Arabidopsis thaliana* development [15]. Disrupted RP genes result in varied, distinctive phenotypes suggesting that not all ribosomal components are equally important all the time. For example, RpL38 mouse mutants exhibit a homeotic transformation phenotype with few other effects [1], whilst RpL38 mutants in *D. melanogaster* exhibit large wings, small bristles, delayed development and disorganised wing hair polarity [16].

Several human diseases, called ribosomopathies, have been attributed to mutations in RP genes. These diseases exhibit varying clinical symptoms between different RP mutations [17, 18]. This suggests human RPs may possess specialised functions, through their requirement for the translation of specific mRNA pools, but this could also be the result of extra-ribosomal RP functions or ribosome insufficiency. Mutations in RpS19, RpS28, RpS10 and RpS5 result in the ribosomopathy Diamond-Blackfan anaemia. Knocking down these RPs specifically results in reduced translation of the erythropoietic transcription factor GATA1, whilst the translation of other mRNAs is unaffected [18]. Therefore, to properly understand ribosomopathies it is necessary to dissect how differences in RP levels impacts ribosome composition and structure.

Human cytoplasmic ribosomes usually comprise of 80 RPs and 4 rRNAs. This is similar across the majority of multicellular eukaryotes, including *D. melanogaster* with 80 RPs and 5 rRNAs. However, there are 93 cytoplasmic RP genes annotated in FlyBase: 39 small subunit proteins and 54 large subunit proteins [19]. These additional genes code for 13 paralogs in *D. melanogaster*. In fact, many RP genes possess paralogs across eukaryotes, for example human RpL3 and RpL3L [13] and *A. thaliana* RpS8A and RpS8B [15]. In total, there are 19 pairs of paralogs in humans [13] and all 80 RPs in *A. thaliana* have paralogs [20]. Expression analyses at RNA and protein levels have indicated there are differences in levels of both canonical RPs and RP paralogs across tissues [21, 22]. But given the possibility for extra-ribosomal effects of these differences, the impact on ribosomes cannot be determined without detailed characterisation of ribosomal composition and structure in concert.

To dissect the function of ribosome heterogeneity it is necessary to understand the biological importance within the context of whole organisms and their development. Within the developmental biology field, a large proportion of research on gene expression control focuses on the contribution of transcription. However, during development a variety of processes and key time points have been shown to be highly dependent on the regulation of mRNA translation, e.g. oogenesis in *Xenopus* [23], early embryo development in *Drosophila* [24] and mammalian erythropoiesis [25]. The balance between self-renewal and differentiation at the stem cell niche is highly dependent on translation in both the ovary and the testis [26]. This is exemplified by disruptions to the stem cell niche in the testis when RPs are knocked down e.g. RpL19 RNAi results in the over-proliferation of early germ cells in *D. melanogaster* [27]. During the meiotic phase of gametogenesis, transcription does not occur, therefore meiotic cells rely on post-transcriptional gene regulation [28]. The translational machinery has evolved to become specialised within the testis with various testis-specific components, e.g. eIF4E-3 in *D. melanogaster* [29]. Many of the RP mutants associated with the *Minute* phenotypes have impaired fertility in both males and females [30, 31]. To date, a robust assessment of ribosome composition across tissues, including gonads, is missing.

Here we hypothesise that specialised ribosomes exist in the *D. melanogaster* testis, to provide an additional level of mRNA translational regulation during spermatogenesis. Thus, we set out to determine potential changes to the *D. melanogaster* ribosome by probing the protein composition in 3 adult tissues (head, testis and ovary) and in embryos. Using quantitative mass spectrometry, we identified heterogeneous ribosome populations, especially in the gonads. The main sources of this variation in ribosome composition are paralog enrichment and paralog-switching, as evidenced by western blotting, cryo-electron microscopy (cryo-EM) and RNA-Seq analysis. We found little difference in composition between single 80S ribosomes and the more translationally active polysome ribosomes from the same tissue. We solved structures of different ribosome populations to understand the potential mechanistic impact of these paralog-switching events, and found structural differences between testis and ovary ribosomes. To understand the broader importance of specialisation through paralog-switching events we analysed the levels of conservation between paralog pairs. We found that RpL22 has a duplicate RpL22L in mammals (including humans), and RpL22-like in *Drosophila*. These duplication events have occurred independently suggesting that it may represent a common mechanism of specialisation across a range of organisms and ribosomes.

## MATERIALS AND METHODS

### Growth conditions

*Drosophila melanogaster* wild type (Dahomey) were raised on standard sugar–yeast agar [32]. Flies were kept at 25°C and 50% humidity with a 12:12 hr light:dark cycle in 6 oz Square Bottom Bottles (Flystuff).

### Tissue harvest

∼300 pairs of ovaries per replicate were harvested from 3-6 day old females in 1X PBS (Lonza) with 1 mM DTT (Sigma) and 1 U/µL RNAsin Plus (Promega) and flash frozen in liquid nitrogen. ∼500 (replicate 1) and ∼1000 (replicates 2 and 3) pairs of testes were harvested from 1-4 day old males in 1X PBS with 2 mM DTT and 1 U/µL RNAsin Plus and flash frozen in groups of ∼10 pairs. ∼500 heads (∼50:50 female:male, 0-4 day old) per replicate were isolated by flash freezing whole flies and subjecting them to mechanical shock to detach heads. Heads were passed through 1 mm mesh filter with liquid nitrogen and transferred to Dounce homogeniser for lysis. ∼500 µL of 0-2 hr embryos per replicate were obtained from cages after pre-clearing for 2 hrs. Laying plates comprised of 3.3% agar, 37.5% medium red grape juice compound (Young’s Brew) and 0.3% methyl 4-hydroxybenzoate, supplemented with yeast paste of active dried yeast (DCL) and dH20. Embryos were washed in dH20 and embryo wash buffer (102.5 mM NaCl (Sigma), 0.04% TritonX-100 (Sigma)) and then flash frozen with minimal liquid.

### Ribosome purification

All stages were performed on ice or at 4°C wherever possible. Ovaries and testes were disrupted using RNase-free 1.5 mL pestles (SLS) in lysis buffer A (50 mM Tris-HCl pH 8 (Sigma), 150 mM NaCl, 10 mM MgCl2 (Fluka), 1% IGEPAL CA-630 (Sigma), 1 mM DTT, 100 µg/mL cycloheximide, 2 U/µL Turbo DNase (Thermo Fisher), 0.2 U/µL RNasin Plus, 1X EDTA-free protease inhibitor cocktail (Roche)). Lysis buffer A does not disrupt embryos present within the ovary sample, since bleach would be required to remove the chorion. Ovaries and testes were lysed in 500 µL lysis buffer A. Heads were lysed using 8 mL Dounce homogeniser with loose pestle in 1.5 mL lysis buffer B (10 mM Tris-HCl pH 7.5 (Gibco), 150 mM NaCl, 10 mM MgCl_2_, 1% IGEPAL CA-630, 1% Triton X-100, 0.5% sodium deoxycholate (Sigma), 2 mM DTT, 200 µg/mL cycloheximide, 2 U/µL Turbo DNase, 40 U/mL RNAsin Plus, 1X EDTA-free protease inhibitor cocktail). Then 500 µL aliquots were transferred to 2 ml Dounce with tight pestle and further lysed for approximately 30 strokes. Embryos were ground in liquid nitrogen using pestle and mortar, and added to lysis buffer B. All lysates were lysed for ≥30 minutes with occasional agitation, then centrifuged for 5 minutes at 17,000 x g to remove cell debris. Head and embryo cytoplasmic supernatants were obtained by avoiding both floating fat and insoluble pellet and repeatedly centrifuged until free of debris.

Cytoplasmic lysates were loaded onto 18 – 60% sucrose gradients (50 mM Tris-HCl pH 8.0, 150 mM NaCl, 10 mM MgCl_2_, 100 µg/mL cycloheximide, 1 mM DTT, 1X EDTA-free protease inhibitor cocktail) and ultra-centrifuged in SW40Ti rotor (Beckman) for 3.5 h at 170,920 x g at 4°C. Ovary and embryo samples were split across two gradients. Fractions were collected using a Gradient Station (Biocomp) equipped with a fraction collector (Gilson) and Econo UV monitor (BioRad). Fractions containing 80S were combined, and same with polysomes (Sup 1A-D). Fractions were concentrated using 30 kDa column (Amicon Ultra-4 or Ultra-15) at 4°C and buffer exchanged (50 mM Tris-HCl pH 8, 150 mM NaCl, 10 mM MgCl_2_) until final sucrose ≥0.1%. Samples were quantified using Qubit Protein Assay Kit (Invitrogen).

For EDTA treatment experiments polysomes were disrupted by the addition of 30 mM EDTA to the lysis buffer and sucrose gradient. MgCl_2_ was also omitted from the lysis buffer and sucrose gradient.

### TMT Labelling and High pH reversed-phase chromatography

An equal amount (TMT1=40 µg, TMT=40 µg, TMT3=35 µg) of each sample was digested with trypsin (2.5 µg trypsin per 100 µg protein; 37°C, overnight), labelled with Tandem Mass Tag (TMT) six or ten plex reagents according to the manufacturer’s protocol (Thermo Fisher Scientific) and the labelled samples pooled.

100 µg aliquots of pooled samples were evaporated to dryness, resuspended in 5% formic acid and then desalted using a SepPak cartridge according to the manufacturer’s instructions (Waters). Eluates from the SepPak cartridge was again evaporated to dryness and resuspended in buffer C (20 mM ammonium hydroxide, pH 10) prior to fractionation by high pH reversed-phase chromatography using an Ultimate 3000 liquid chromatography system (Thermo Scientific). In brief, samples were loaded onto an XBridge BEH C18 Column (130Å, 3.5 µm, 2.1 mm X 150 mm, Waters) in buffer C and peptides eluted with increasing gradient of buffer D (20 mM Ammonium Hydroxide in acetonitrile, pH 10) from 0-95% over 60 minutes. The resulting fractions were evaporated to dryness and resuspended in 1% formic acid prior to analysis by nano-LC MS/MS using an Orbitrap Fusion Tribrid mass spectrometer (Thermo Scientific).

### Nano-LC Mass Spectrometry

High pH RP fractions were further fractionated using an Ultimate 3000 nano-LC system in line with an Orbitrap Fusion Tribrid mass spectrometer (Thermo Scientific). In brief, peptides in 1% (vol/vol) formic acid were injected onto an Acclaim PepMap C18 nano-trap column (Thermo Scientific). After washing with 0.5% (vol/vol) acetonitrile 0.1% (vol/vol) formic acid peptides were resolved on a 250 mm × 75 μm Acclaim PepMap C18 reverse phase analytical column (Thermo Scientific) over a 150 minute organic gradient, using 7 gradient segments (1-6% solvent B over 1 minute, 6-15% B over 58 minutes, 15-32% solvent B over 58 minutes, 32-40% solvent B over 5 minutes, 40-90% solvent B over 1 minute, held at 90% solvent B for 6 minutes and then reduced to 1% solvent B over 1 minute) with a flow rate of 300 nl minutes^−1^. Solvent B was aqueous 80% acetonitrile in 0.1% formic acid. Peptides were ionized by nano-electrospray ionization at 2.0 kV using a stainless-steel emitter with an internal diameter of 30 μm (Thermo Scientific) and a capillary temperature of 275°C.

All spectra were acquired using an Orbitrap Fusion Tribrid mass spectrometer controlled by Xcalibur 2.1 software (Thermo Scientific) and operated in data-dependent acquisition mode using an SPS-MS3 workflow. FTMS1 spectra were collected at a resolution of 120 000 with an automatic gain control (AGC) target of 200 000 and a max injection time of 50 ms. Precursors were filtered with an intensity threshold of 5000, according to charge state (to include charge states 2-7) and with monoisotopic peak determination set to peptide. Previously interrogated precursors were excluded using a dynamic window (60 s +/-10 ppm). The MS2 precursors were isolated with a quadrupole isolation window of 1.2 m/z. ITMS2 spectra were collected with an AGC target of 10 000, max injection time of 70 ms and CID collision energy of 35%.

For FTMS3 analysis, the Orbitrap was operated at 50 000 resolution with an AGC target of 50 000 and a max injection time of 105 ms. Precursors were fragmented by high energy collision dissociation (HCD) at a normalised collision energy of 60% to ensure maximal TMT reporter ion yield. Synchronous Precursor Selection (SPS) was enabled to include up to 5 MS2 fragment ions in the FTMS3 scan.

### TMT Data Analysis

The raw data files were processed and quantified using Proteome Discoverer software v2.1 (Thermo Scientific) and searched against the UniProt *Drosophila melanogaster* database (downloaded March 2018; 41,157 sequences) using the SEQUEST HT algorithm. Peptide precursor mass tolerance was set at 10 ppm, and MS/MS tolerance was set at 0.6 Da. Search criteria included oxidation of methionine (+15.995 Da) and acetylation of the protein N-terminus (+42.011 Da) as variable modifications and carbamidomethylation of cysteine (+57.021 Da) and the addition of the TMT mass tag (+229.163 Da) to peptide N-termini and lysine as fixed modifications. Searches were performed with full tryptic digestion and a maximum of 2 missed cleavages were allowed. The reverse database search option was enabled and all data was filtered to satisfy false discovery rate (FDR) of 5% [33] and reported.

Peptide IDs not corresponding to *D. melanogaster* proteins were removed from all TMT replicates. Using the protein grouping decided by PD2.1, master protein selection was improved using an in-house script to select the UniProt accession (database downloaded Jan 2021; 42,818 sequences) with the best annotation whilst maintaining confidence in protein identification and quantitation. This resulted in a list of 1906 proteins for TMT1, 3613 proteins for TMT2 and 1869 proteins for TMT3. Abundances are the sum of the S/N values for the TMT reporter groups for all peptide-spectrum matches (PSMs)matched to the protein. Normalised abundances of these values were obtained by normalising the Total Peptide Amount in each sample such that the total signal from each TMT tag is the same. Scaled abundances are either normalised abundances scaled to a pooled sample or the normalised abundance scaled to the average of all samples within that replicate. Data from all three replicates was merged into one data sheet from which comparisons between tissues were made with statistical analyses for all proteins detected (Sup Table 1).

Pair-wise comparisons were made to calculate the log_2_ fold-change difference of proteins across different tissue samples. For TMT1 and 3, normalised abundances were used, and for TMT2 the normalised abundance was scaled to a pooled sample to allow comparison between TMT runs. Standard t-test was used to test the statistical significance of the log_2_ fold-changes between tissue samples. Analysis of TMT data and hierarchical clustering were performed in R.

Fold-change differences between highly similar peptides were determined to estimate relative abundances of different RP paralog pairs between tissue samples. PSMs were only used for this purpose if the peptides were of identical length and had fewer than 2 amino acid changes.

### Antibodies and western blotting

RP paralog specific antibodies were generated using custom peptides for RpL22 (CNKGDTKTAAAKPAEK), RpL22-like (CSSQTQKKNASKAKSK), RpS19a (CQIVFKQRDAAKQTGP), RpS19b (CKQRERSAPVSMIITT), RpL37a (CREGTQAKPKKAVASK) and RpL37b (CRNGLREGGAAKKKTN) (Pepceuticals, UK). RpL22 and RpL22-like antibodies were isolated from serum via affinity purification using HiTrap NHS-activated HP columns (Cytiva Life Sciences). RpS5a and RpS5b antibodies were kindly gifted by the Lasko lab [34].

Protein samples were separated on a 4-20% Mini-Protean TGX gel before being transferred to 0.2 μm nitrocellulose membrane, which was blocked for 1h in 5% milk in 1X TBST. Membranes were probed with antibodies diluted in 1X TBST. Primary antibody concentrations were: RpL22 1:2500, RpL22-like 1:2500, RpS5a 1:1000, RpS5b 1:1000, RpS19a 1:25, RpS19b 1:25, RpL37a 1:25, RpL37b 1:25, RpL40 1:1000 (ab109227, abcam).

### Source of RNA-Seq data

RNA-Seq data was extracted from ModMine (intermine.modencode.org)[35] with data from modENCODE project [36, 37]. Values are Reads Per Kilobase of transcript, per Million mapped reads (RPKMs).

### Cryo-EM

400 mesh copper grids with a supporting carbon lacey film coated with an ultra-thin carbon support film < 3 nm thick (Agar Scientific, UK) were employed. Grids were glow-discharged for 30 seconds (easiGlow, Ted Pella) prior to applying 3 µL of purified ribosomes, and vitrification was performed by plunge-freezing in liquid ethane cooled by liquid nitrogen using a Leica EM GP device (Leica Microsystems). Samples were diluted using the buffer exchange buffer (50 mM Tris-HCl pH 8, 150 mM NaCl, 10 mM MgCl_2_) as required. Cryo-EM data was collected on a FEI Titan Krios (Astbury Biostructure Laboratory, University of Leeds) EM at 300 kV, using a total electron dose of 80 e^-^/Å^2^ and a magnification of 75,000 × at -2 to -4 μm defocus. Movies were recorded using the EPU automated acquisition software on a FEI Falcon III direct electron detector, in linear mode, with a final pixel size of 1.065 Å/pixel (Sup Table 2).

### Image processing

Initial pre-processing and on-the-fly analysis of data was performed as previously described [38]. Image processing was carried out using RELION 2.0/2.1 or 3.0 [39]. MOTIONCOR2 [40] was used to correct for beam-induced motion and calculate averages of each movie. gCTF [41]. was used for contrast transfer function determination. Particles were automatically picked using the Laplacian of Gaussian function from RELION [42]. Particles were classified using two rounds of reference-free 2D classification. For the testis 80S reconstruction, particles contributing to the best 2D class averages were then used to generate an initial 3D model. For the ovary 80S and testis polysome reconstruction, the testis 80S average was used as initial reference. Particles were classified by two rounds of 3D classification, and the best 3D classes/class were 3D refined, followed by per-particle CTF correction and Bayesian polishing [42]. Post-processing was employed to mask the model, and to estimate and correct for the B-factor of the maps [43]. The testis 80S map was further processed by multi-body refinement, as previously described [44]. The final resolutions were determined using the ‘gold standard’ Fourier shell correlation (FSC = 0.143) criterion (Sup Table 2). Local resolution was estimated using the local resolution feature in RELION.

### Atomic modelling

The *D. melanogaster* embryo ribosome (PDB code 4V6W) was used as a model to calculate the structures of the testis and ovary ribosomes. First, the full atomic model was fitted into the testis 80S cryo-EM average using the ‘fit in map’ tool from Chimera [45]. Then, fitting was refined by rigid-body fitting individual protein and RNA PDBs into the maps using Chimera. The 18S and 28S ribosomal RNAs were split into two separate rigid bodies each. Proteins and RNAs not present in our averages (i.e. elongation factor 2 and Vig2 for all models, and E-tRNA) and proteins and RNA with poor densities (i.e. RpLP0 and RpL12, and some regions of the 18S and 28S ribosomal RNAs) were removed at this stage. The paralog proteins used for each ribosome are listed in Table 1. For the testis 80S atomic model, IFRD1 was modelled using SWISS-MODEL [46], based on the atomic model for rabbit IFRD2 (PDB model 6MTC). For the testis polysome model, the mRNA was based on PDB model 6HCJ, and the P-tRNA on the E-tRNA from PDB model 4V6W. The full atomic models were refined using Phenix [47], and the paralogs listed in Fig 2A (plus RpL31 and RpS18) were manually inspected and corrected using COOT [48] (except Rp10Ab, which was not manually inspected due to the low resolution of that area in the average maps, and RpLP0, which was not present in the model). This cycle was repeated at least three times per ribosome model. The quality of the atomic models was assessed using the validation server from the PDB website (https://validate-pdbe.wwpdb.org/). As the 60S acidic ribosomal protein P0 deposited (RpLP0) in the PDB (4V6W) is from *Homo sapiens*, we generated a *D. melanogaster* homology model using SWISS-MODEL. This protein was rigid-body fitted using Chimera after the atomic model refinement and is displayed in Fig 4 for relative position and size comparison purposes only. Figures were generated using Chimera.

**Table 1:** Atomic model paralog comparison. Switched paralogs are highlighted in bold. L31 and S18 do not have paralogs and were used as internal controls. Resolution refers to the local resolution of the testis 80S EM average for each paralog. The % identity refers to identity when comparing the amino acids included in the refinement. RMSD is Root-mean-square deviation in Å between the atomic models.

|  |  | Testis 80S |  |  | Ovary 80S |  |  |  |
| --- | --- | --- | --- | --- | --- | --- | --- | --- |
|  |  | Paralog in Testis | # aac included in refinement | Local Resolution (Å) | Paralog in ovary | # aac included in refinement | % identity | RMSD across all atom pairs (Å) |
| Large subunit | RpL31 | RpL31 | 108 | 4 | RpL31 | 108 | 100 | 0.616 |
|  | RpL7/L7-like | RpL7 | 223 | 3.5 | RpL7 | 226 | 100 | 0.399 |
|  | RpL34a/b | RpL34b | 103 | 3.5 | RpL34b | 103 | 100 | 0.460 |
|  | RpL24/L24-like | RpL24 | 58 | 3.5 | RpL24 | 60 | 100 | 0.518 |
|  | **RpL37a/b** | **RpL37b** | **87** | **3** | **RpL37a** | **87** | **76** | **0.714** |
|  | **RpL22/L22-like** | **RpL22-like** | **96** | **4.5** | **RpL22** | **99** | **58** | **3.078** |
| Small subunit | RpS18 | RpS18 | 134 | 5 | RpS18 | 134 | 100 | 1.591 |
|  | RpS15Aa/b | RpS15Aa | 129 | 3.5 | RpS15Aa | 129 | 100 | 0.547 |
|  | **RpS5a/b** | **RpS5a** | **189** | **5.5** | **RpS5b** | **189** | **90** | **1.852** |
|  | RpS28a/b | RpS28b | 62 | 5.5 | RpS28b | 62 | 100 | 1.884 |
|  | RpS19a/b | RpS19a | 139 | 6 | RpS19a | 143 | 100 | 2.613 |
|  | RpS10a/b | RpS10b | 90 | 6.5 | RpS10b | 90 | 100 | 2.717 |

### Vertebrate dataset construction

Coding DNA sequence (CDS) data for 207 vertebrate animals and 4 non-vertebrates (*D. melanogaster*, two *Caenorhabditis* species and *S. cerevisiae*) were obtained from Ensembl (release 97, [49]). We performed homology searches using two human RpL22 family proteins (RPL22 and RPL22L) against 6,922,005 protein sequences using BLASTp (e^-5^) [50]. We identified 1,082 potential RpL22 proteins from 185 vertebrates and 4 non-vertebrates, which were homologous to one or both human RpL22 proteins. As an initial step to reduce the amount of redundancy in the vertebrate dataset, 181 potential RpL22 proteins from 42 selected vertebrates (including humans) were retained to represent as broad a taxonomic sampling of the group. All non-vertebrate sequences, with the exception of two *S. cerevisiae* RpL22 proteins (RPL22A and RPL22B), were also removed from the dataset. 92 alternative transcripts and spurious hits were removed from the dataset through manual cross-validation with Ensembl Genome Browser to give a total of 87 vertebrate and 2 yeast RpL22 family proteins.

### Invertebrate dataset construction

A CDS dataset for 78 invertebrate animals was obtained from Ensembl Metazoa (release 44, [49]). The sequence homology search was performed using two *D. melanogaster* RpL22 family proteins (RPL22 and RPL22-like) were searched against 1,618,385 protein sequences using BLASTp (e^-5^) [50]. BLASTp identified 90 potential RpL22 family proteins across 70 invertebrates, which were homologous to one or both *D. melanogaster* RpL22 proteins. 15 alternative transcripts and spurious hits were removed from the dataset through manual cross-validation with Ensembl Genome Browser to give a total of 75 invertebrate RpL22 family proteins. Together with 87 vertebrate and 2 outgroup proteins, our final dataset consisted of 164 RpL22 family proteins sampled across the metazoan tree of life.

### Phylogenetic reconstructions of metazoan RpL22 family

Initial phylogenetic reconstruction of the metazoan RpL22 family was performed using the full dataset of 164 sequences (87 invertebrate sequences, 75 vertebrate sequences and two yeast sequences). All sequences were aligned using three different alignment algorithms: MUSCLE [51], MAFFT [52] and PRANK [53]. MUSCLE was run with the default parameters, and MAFFT was run with the automatically-selected most-appropriate alignment strategy (in this case, L-INS-I). PRANK was run with both the default parameters and the PRANK+F method with “permanent” insertions. All four resultant alignments were compared against each other using MetAl [54], and were all judged to be mutually discordant based on differences of 20-25% between each pair of alignments. Column-based similarity scores were calculated for each alignment using the norMD statistic [55]. The MUSCLE alignment had the highest column-based similarity score (1.281) and was selected for further analysis. This alignment was trimmed using TrimAl’s gappyout method [56]. Maximum-likelihood phylogenetic reconstruction was performed on the trimmed alignment using IQTREE [57], with a WAG+R6 model selected by ModelFinder Plus [58] and 100 bootstrap replicates.

A smaller, taxonomically-representative RpL22 family dataset containing 50 RpL22 genes from 30 animals and *S. cerevisiae* was constructed for a representative RpL22 family phylogeny. This dataset was aligned using the same four methods described above, and all alignments were judged to be mutually discordant (differences of 19-37%) using MetAl [54]. The MUSCLE alignment had the highest column-based similarity score assigned by norMD (0.702) and was selected for further analysis. As above, this alignment was trimmed using TrimAl’s gappyout method. Maximum-likelihood phylogenetic reconstruction was performed on the trimmed alignment using IQTREE [57], with a DCMut+R3 model selected by ModelFinder Plus [58] and 100 bootstrap replicates.

### Data availability

The mass spectrometry proteomics data have been deposited to the ProteomeXchange Consortium via PRIDE partner repository with the dataset identified PDX026227. The EM-density maps for testis 80S, testis polysomes and ovary 80S are deposited in the EMDB under the accession numbers EMD-10622, EMD-10623 and EMD-10624. The refined models are deposited in the PDB under accession codes 6XU6, 6XU7and 6XU8.

## RESULTS

### Heterogeneous ribosome populations exist in different tissues

Many eukaryotic genomes contain numerous RP paralogs yet their contribution to ribosomal function is poorly understood. In *D. melanogaster* there are 93 RP genes (FlyBase), which include 13 pairs of paralogs, resulting in 80 proteins in each ribosome [59]. The expression of RPs and specifically RP paralogs has been reported to vary in a tissue specific manner [15, 21]. To profile potential differences in expression in *D. melanogaster* we analysed publicly available RNA-Seq data across various developmental time points and tissues. Hierarchical clustering of RP mRNA abundances across these different biological samples reveals variations in expression of RP mRNAs between tissues, with a cluster of RPs with much higher expression in the testis compared to other tissues (Fig 1A). This includes RpL22-like, a paralog of RpL22 previously reported as a testis-specific ribosomal protein [7]. This result suggests the presence of testis-specific translational machinery.

**Figure 1:**
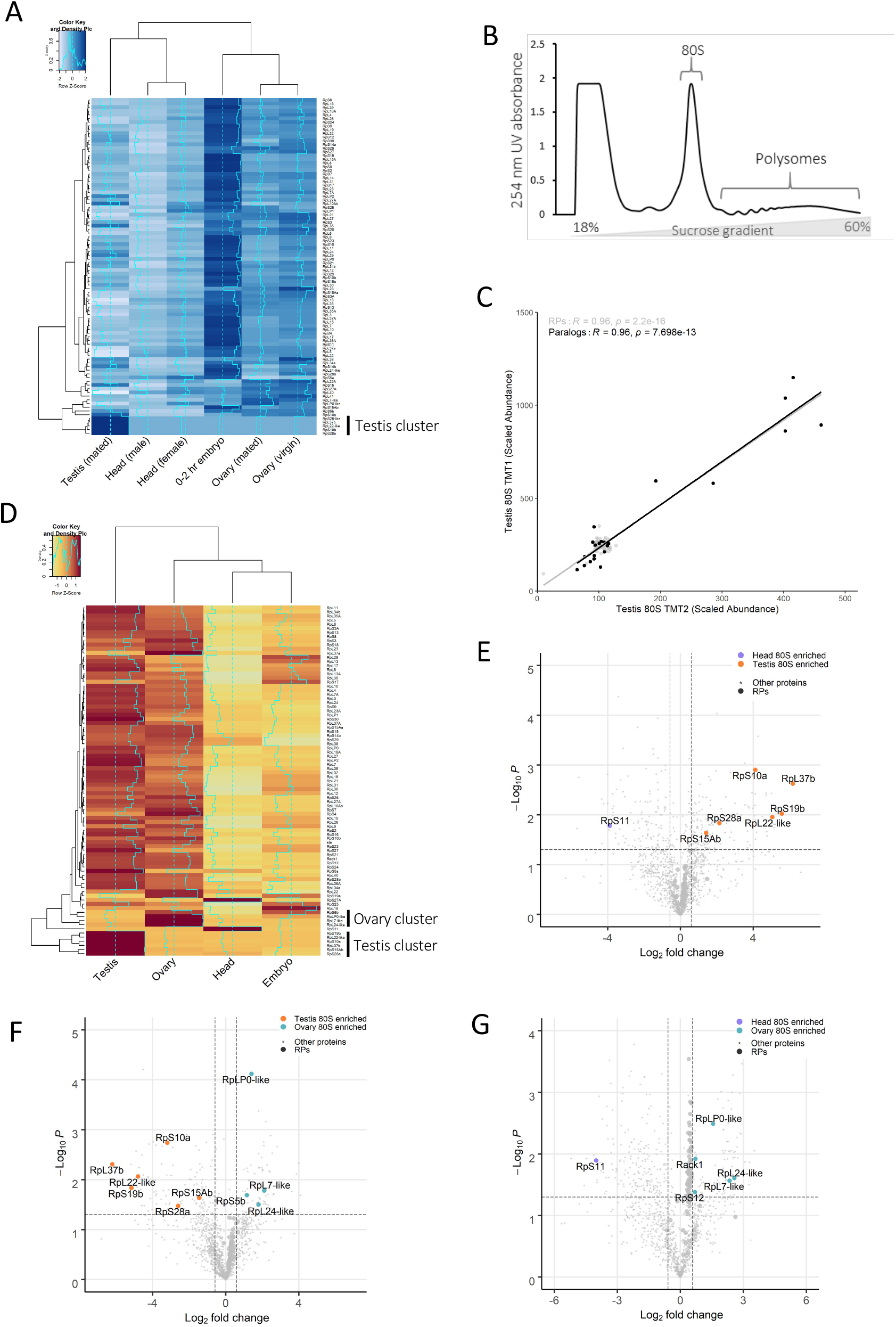
Heterogeneous ribosome populations exist in different tissues. (A) Hierarchical clustering of modENCODE RNA-Seq data (RPKM) for ribosomal proteins (RPs) across testis, ovary, head and embryo reveals differences in RP expression clustered according to row i.e. ribosomal protein with Z-scores calculated and plotted. The analysis shows a cluster of 5 testis-specific RPs. (B) Schematic of strategy used to isolate and compare 80S ribosome and polysome complex composition. (C) Correlation of two biological repeats of TMT mass spectrometry experiments for scaled protein abundances within 80S ribosomes isolated from testis shows replicates are reproducible. Correlations are shown for all RPs (grey) and RP paralogs (black) with Pearson’s correlation coefficients calculated. This shows replicates are reproducible. (D) Hierarchical clustering of log_2_ scaled protein abundances from replicate 2. Normalised abundances were scaled to a control pool from which Z-scores were calculated and plotted. Heatmap is clustered according to row i.e. ribosomal protein. (E-G) Volcano plots highlighting the differences in ribosomal protein composition between (E) head and testis, (F) testis and ovary, and (G) ovary and head tissues. Enriched proteins have a >1.5-fold change with a p-value of <0.05. Enriched RPs are labelled.

To determine whether these different RPs are translated and incorporated into ribosomes we assessed the protein composition of ribosomes from testes, ovaries, heads (mixture of male and female) and embryos (0-2 hr). Ribosomal complexes were purified using sucrose gradients and ultracentrifugation (Fig 1B). Both 80S (monosome) and polysome complexes were isolated. The relative amounts of ribosomes existing as 80S or polysome complexes varied substantially across the samples (Sup 1). In general, polysome levels are substantially lower than monosome levels in tissues (Sup 1), as previously shown *in vivo* [60], especially when compared to cultured cells [61]. Both monosome and polysome fractions were isolated for each tissue and subjected to quantitative mass spectrometry (tandem mass tag; TMT). Overall correlation between the biological replicates is high as RPs in testis 80S samples had Pearson’s correlation coefficients of 0.96, 0.95 and 0.96 (Fig 1C, Sup 2). Similar results are obtained when considering only ribosomal paralogs and across samples (Sup 2). We only used unique peptides in our analysis, which is particularly important when distinguishing between paralogs (Sup 3 and 4).

To understand differences in ribosome composition in 80S complexes between the tissues, protein abundances of RPs were subject to hierarchical clustering (Fig 1D). Two clear protein clusters emerged. One where proteins are enriched in the testis 80S ribosomes compared to 80S ribosomes from other tissues, including RpL22-like, RpL37b, RpS19b, RpS10a and RpS28a, RpS15Ab. The other is an ovary 80S enriched cluster of ribosomal proteins, including RpS5b, RpL24-like, RpL7-like and RpL0-like (Fig 1D). Similar clusters are seen for the three biological repeats (Fig 1D, Sup 5A and B). To identify substantial and statistically significant differences in ribosomal protein composition, differences in abundances were plotted between different 80S complexes, employing 1.5 fold-change and 0.05 p-value as cut-offs. Comparison of testis 80S with head 80S and ovary 80S revealed that the same 6 RPs (RpL22-like, RpL37b, RpS19b, RpS10a, RpS28a and RpS15Ab) are highly enriched in the testis 80S (Fig 1E and F). Likewise, comparison of ovary 80S with testis 80S and head 80S revealed that the same 4 RPs (RpS5b, RpL24-like, RpL7-like and RpL0-like) are enriched in ovary 80S (Fig 1F and G). Additionally, the comparison between ovary 80S with head 80S revealed that RpS12 and RACK1 are also enriched in ovary 80S compared to head 80S. The comparisons involving head 80S showed an enrichment of RpS11 in the head 80S (Figs 1E and G). Embryo 80S showed no paralog enrichment (Sup 5C-E). Overall, heterogeneity seems most common in the gonads and we identify both testis- and ovary-enriched RPs.

### Ribosomal protein paralogs contribute to ribosome heterogeneity

There are 13 pairs of RP paralogs in the *D. melanogaster* genome and from our TMT data we can see the majority are both expressed and incorporated into 80S ribosomes in at least one of the analysed tissues. In fact, out of the enriched RPs we identified, 10 out of 13 were paralogs (Figs 1E-G). Hierarchical clustering of just RP paralogs re-emphasises the existence of gonad specific ribosomal complexes mostly through changes to RP paralogs (Fig 2A). To understand the relationship between each of the two paralogs we compared the quantitative differences in abundances between tissues for each paralog pair (i.e. log_2_ fold-change – log_2_ FC – between ovary and testis, Fig 2B). RpS14a and RpS14b have the same amino acid sequences, so are indistinguishable by mass spectrometry and therefore excluded from this analysis. In addition, RpL10Aa was also omitted because no unique peptides were detected. Calculating log_2_ FC between ovary and testis 80S complexes indicates that out of 11 pairs we could analyse in this way, the 6 testis-specific paralogs are enriched in the testis and the 4 ovary-specific paralogs are enriched in the ovary. Only for the RpL34 paralog pair did neither paralog show differential incorporation into the 80S ribosome between ovary and testis. Overall, only one paralog from each pair is significantly enriched in either ovary or testis, whilst the other paralog in each pair shows little or no difference (i.e. log_2_ FC below 0.5). In general, these other paralogs are slightly enriched in the other tissue but not significantly. For example, for the paralog pair RpS5, RpS5b is enriched in the ovary (*p* value = 0.02) and RpS5a is more enriched in the testis (*p* value = 0.026).

**Figure 2:**
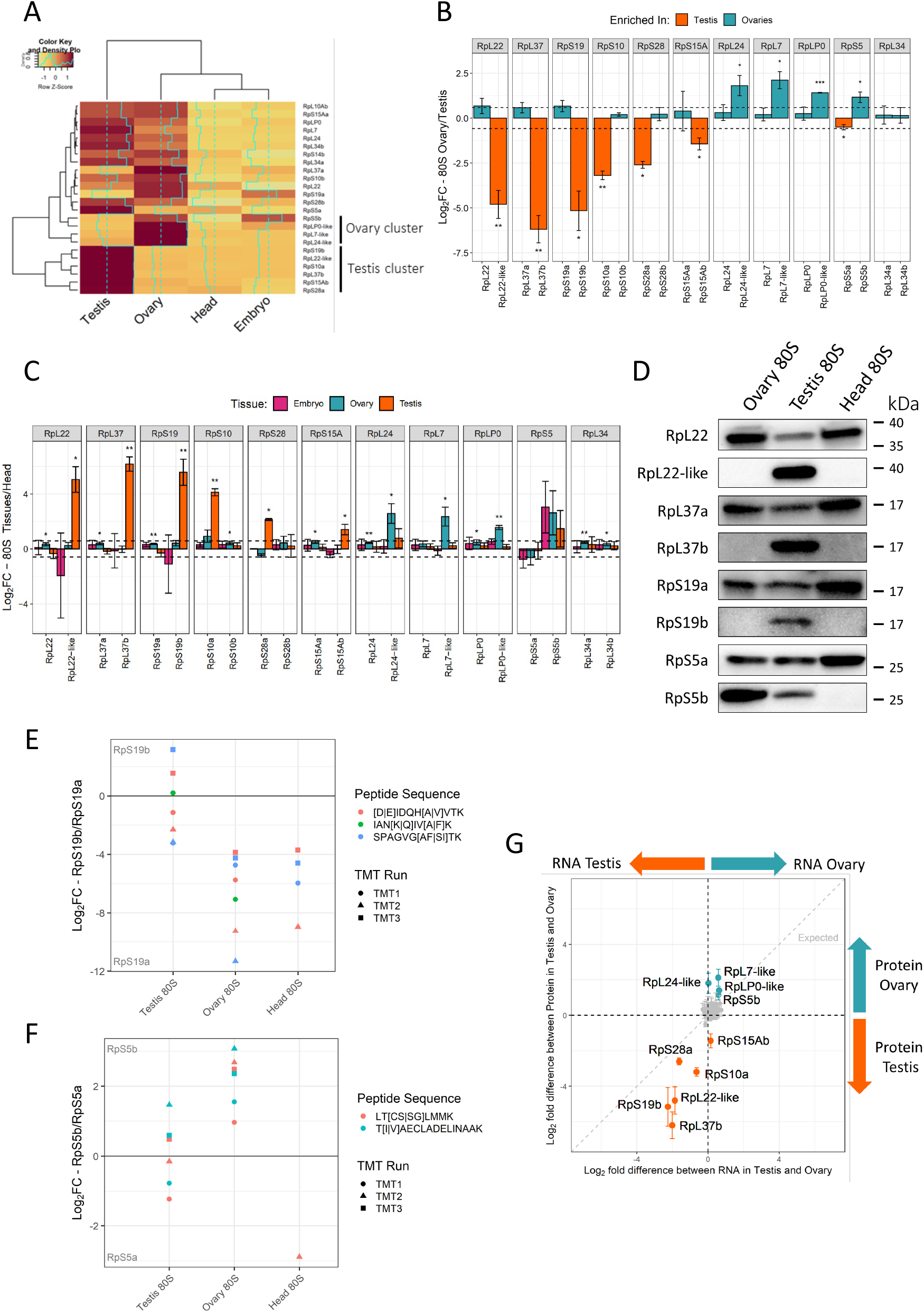
Gonad ribosome heterogeneity through paralog enrichment and paralog-switching. (A) Hierarchical clustering of log_2_ scaled abundances for the 24 RP paralogs across the 4 tissues from replicate 2, scaled by row. (B) Comparison of paralog pairs in testis and ovary 80S ribosomes. Bars represent log_2_ fold-change between the two tissues. Dotted lines represent a log_2_ fold-change of 1.5, p-value <0.05, ** <0.01, *** <0.001. (C) Comparison of paralog pairs in all tissues relative to head tissue. Dotted lines represent a log_2_ fold-change of 1.5, * p-value <0.05, ** <0.01, *** <0.001. (D) Immunoblots of purified 80S monosomes from ovary, testis and heads tissues using paralog specific antibodies. (E and F) Comparison of highly similar unique peptides for (E) RpS19a/b and (F) RpS5a/b in testis, ovary, and head tissues. Peptides compared for each paralog pair are of the same size and have 1 or 2 amino acid changes (differences shown in the key within square brackets). Comparisons are shown as log_2_ fold-change differences for each paralog pair. (G) Log_2_-fold difference plot comparing testis and ovary at both RNA (ovary RNA, mated females) and protein levels. All 85 RPs detected in our TMT experiments are plotted with testis/ovary specific paralogs labelled. Error bars represent standard deviations. Grey dotted line represents expected relationship between fold-change in RNA and protein.

To further compare across all the tissues in this way we calculated Log_2_ fold-change of embryo, ovary and testis to head, because head showed little heterogeneity (Fig 2C). This analysis shows the same testis-specific and ovary-specific paralogs (except for RpS5, which is not significant in this analysis). Interestingly RpS5b is enriched in embryo, ovary and testis, when compared to the head, but is not statistically significant. Thus, RpS5b has an unusually broad incorporation across the different sampled ribosomes.

To validate the difference in paralog levels seen by TMT we generated paralog specific antibodies for RpL22, RpL22-like, RpL37a, RpL37b, RpS19a and RpS19b. Using these and published RpS5a and RpS5b antibodies [34] we probed paralog levels in purified 80S ribosomes isolated from ovaries, testes and heads. This analysis confirmed that RpL22-like is enriched in the testis when compared to ovary and head 80S ribosomes (Fig 2D). Furthermore, it showed a concomitant reduction of RpL22 levels in testis 80S, the extent of which indicates that a larger proportion of the testis 80S ribosomes contain RpL22-like rather than RpL22. A similar pattern was seen for RpL37b and RpL37a, again suggesting that a larger proportion of the testis 80S ribosomes contain RpL37b rather than RpL37a. RpS19b was also detected in testis 80S ribosomes, but without such a large reduction in RpS19a when compared to ovary and head. This suggests that although RpS19b is enriched in the testis, it is present in a similar proportion of ribosomes than RpS19a. The westerns also confirmed the enrichment of RpS5b in the ovary and testis compared with head. This is accompanied by a concomitant reduction of RpS5a in both tissues. Additionally, there is a larger proportion of ovary 80S ribosomes that contain RpS5b instead of RpS5a.

To provide a more accurate comparison of protein levels between each pair of paralogs a detailed analysis of peptides resulting from the paralogs were performed. Pairs of unique peptides in the TMT data were identified, which corresponded to almost identical parts of the two paralogs but were different by 1 or 2 amino acids. These peptides were different enough that they could be confidently assigned to one of the two paralogs e.g.[D]IDQH[A]VTK from RpS19b and [E]IDQH[V]VTK from RpS19a, but similar enough that the quantification can be directly compared. Only similar peptides with comparable characteristics (e.g. charge and length) can be compared by TMT; therefore we reasoned that highly homologous peptides with only 1 or 2 amino acid changes qualified for this criteria. Differences in levels were calculated for each pair for each replicate. For some paralog pairs there were several such peptides that could be analysed whereas for other paralogs, only 1 peptide fitted these parameters. Additionally, some paralogs did not contain unique peptides that allowed for this analysis; these included RpL22 and RpL22-like; and RpL37a and RpL37b. This analysis revealed that the fold difference between RpS19b and a, is ∼0 in testis, indicating that around equal levels of the two paralogs are present within testis 80S ribosomes (Fig 2E). Whereas in ovary and head there is >16 times more RpS19a than there is RpS19b. This analysis confirmed paralog switching for RpS5b in ovary (Fig 2F), i.e. there is ∼2-8 fold more RpS5b in ovary 80S ribosomes than RpS5a. Furthermore, in the testis 80S ribosomes the fold difference suggests there are near equal quantities of RpS5b and RpS5a present (Fig 2F). We note these results correlate with the western blots for the RpS19a/b and RpS5a/b paralogs pairs (Fig 2D). Similar analysis for RpS15Ab/a indicates that although RpS15Ab is enriched in the testis 80S ribosomes, it still represents a minor part of the population compared to RpS15Aa (Sup 6A). For both RpS10b/a (Sup 6B) and RpS28b/a (Sup 6C) only 1 peptide pair could be used for this analysis, and these peptides were not detected in all replicates, therefore we are unable to draw such clear conclusions.

However, this data does suggest that even though RpS10a and RpS28a are enriched in testis 80S ribosomes they are both present at relatively low levels compared to RpS10b and RpS28b.

In summary these data indicate that there are 6 paralogs are enriched in testes ribosomes and 4 in ovaries; and that some of these alternative paralogs are present in a larger proportion of ribosomes than the canonical paralog (i.e. RpL22-like, RpL37b in testis 80S and RpS5b in ovary 80S). Furthermore, RpS19b and RpS5b are present at similar levels to their canonical paralogs in the testis 80S. Together this suggests that there is RP paralog switching occurring in the gonads, whereby the canonical paralog is switched for an alternative one in the majority of ribosomes.

### Differences in ribosome composition are not simply the result of expression differences

To understand the expression of RP paralogs, we analysed RNA levels of each of the paralog pairs (Sup 7) from published RNA-Seq datasets of the tissues we performed TMT in [35-37]. RNA-Seq levels for several of the gonad-enriched paralogs are similar to, or above levels of the canonical paralog in each pair. For example, this is the case for RpL22-like, RpL37b, RpS19b, RpS10a, RpS5b in the testis, and Rp5b in the ovary (Sup 7). Therefore this RNA-Seq data supports the finding that several paralogs can be considered more than simply testis or ovary-enriched, likely present in a substantial proportion of ribosomes and potentially gonad-switched.

Differences in RNA-Seq levels between testis and ovary were then compared to differences in TMT abundance to examine if paralog enrichment is simply the result of differential expression. An increased RNA expression for a given paralog was generally associated with larger log_2_ fold-change in paralog abundance with the 80S ribosome (e.g. RpS19b and RpS28a for testis; and RpS5b and RpLP0-like for ovaries; Fig 2G). However, it is clear that differences in protein composition of ribosomes are not simply driven by transcriptional control of paralog genes. Specifically, we identified several instances when RP incorporation into the ribosome does not correlate with mRNA expression level (Fig 2G). For example, RpL24-like is transcribed at similar levels in ovary and testis (Sup 7) but RpL24-like is far more abundant in ovary 80S than testis 80S ribosomes (Fig 2G). The opposite is seen for RpS15Ab, whose RNA levels are similarly low between testis and ovary but is preferentially incorporated into testis 80S (Sup 7 and Fig 2G). In general, differences in RP paralog enrichment in tissues are driven by differences in mRNA expression, however, additional regulation is taking place for some RPs, such as RpL24-like and RpS15Ab.

### Composition of 80S ribosomes and polysomal ribosomes is similar

There is conflicting evidence as to the functionality or translational activity of monosomes (80S ribosomes). Some studies suggest that these ribosomes are actively translating [62] whilst others suggest that not all 80S ribosomes are engaged in active translation [63]. To determine if there was any difference in ribosome composition between monosomes and polysome complexes for a given tissue, we compared the two by TMT. In general, there is very little difference in RP composition between 80S ribosomes and polysomes (Sup 8). No differences were found between embryo 80S and embryo polysomes (Sup 8A). However, RpL7-like and RpL24-like are enriched in testis 80S (Sup 8B) and RpL24-like is enriched in head polysomes when compared to head 80S ribosome complexes (Sup 8C). Furthermore, a non-paralog ribosomal protein, RpL38, is enriched in ovary polysomes (Sup 8D). Overall, we found larger differences in protein composition between different tissues, than between 80S and polysomal ribosomes from the same tissues, and no consistent differences between 80S and polysome ribosomes. Together this suggests that from the perspective of ribosomal protein composition, there is little difference between 80S and polysomal ribosome complexes.

### Paralog enrichment is within ribosomal complexes

To ensure that the complexes we have analysed represent ribosomes, rather than other large protein complexes we assessed the sensitivity of ovary ribosomal complexes to EDTA. EDTA chelates Mg^2+^ and therefore causes ribosomes to disassociate into 40S and 60S subunits. The distribution of RPs across sucrose gradients in presence and absence of EDTA was assessed by western blots. These reveal a shift of RPs towards 40S and 60S subunit fractions (Sup 9), suggesting that their presence in the complexes we purified was the result of ribosomes. Specifically, RpL22, which is highly abundant in ovary ribosomes, showed a shift from large complexes in the sucrose gradient (polysomes) to small complexes (including 60S subunits) in the presence of EDTA. This was also the case for the two RpS5 paralogs, of which RpS5b is ovary-switched. Both paralogs shift out of the polysomes to small complexes, corresponding to 40S subunits. The patterns exhibited for paralogs were also seen with a canonical RP, RpL40. Together these results indicate that we have identified changes in ribosome protein composition rather than in large non-ribosomal complexes.

### Cryo-electron microscopy of testis and ovary ribosomes reveals a mechanism for inactivation of testis 80S ribosomes

To understand the molecular implications of the paralog switching events we identified by mass spectrometry and western blot, we sought to solve structures of different ribosome populations. Ribosomal complexes were isolated by sucrose gradient centrifugation, in the same way as for the TMT (Fig 1B). Imaging the sample by cryo-electron microscopy (cryo-EM) confirmed that the ribosome complexes were highly pure and concentrated (Sup 10A) and a dataset containing ∼47,000 particles was collected. Three-dimensional classification of this testis 80S dataset identified a single structurally distinct class of 80S ribosomes, which was refined to an average 3.5 Å resolution (Fig 3A and Sup 10B and C). This provided a substantial improvement to the only other *D. melanogaster* ribosome cryo-EM average at 6 Å resolution, from embryos [59]. We performed a similar experiment with ovary 80S ribosome preparations, collecting a dataset containing ∼200,000 particles, and resulting in an average 3.0 Å resolution (Fig 3B; Sup 10D-F). These averages allowed us to generate atomic models for testis and ovary 80S ribosomal complexes (Sup Table 2).

**Figure 3:**
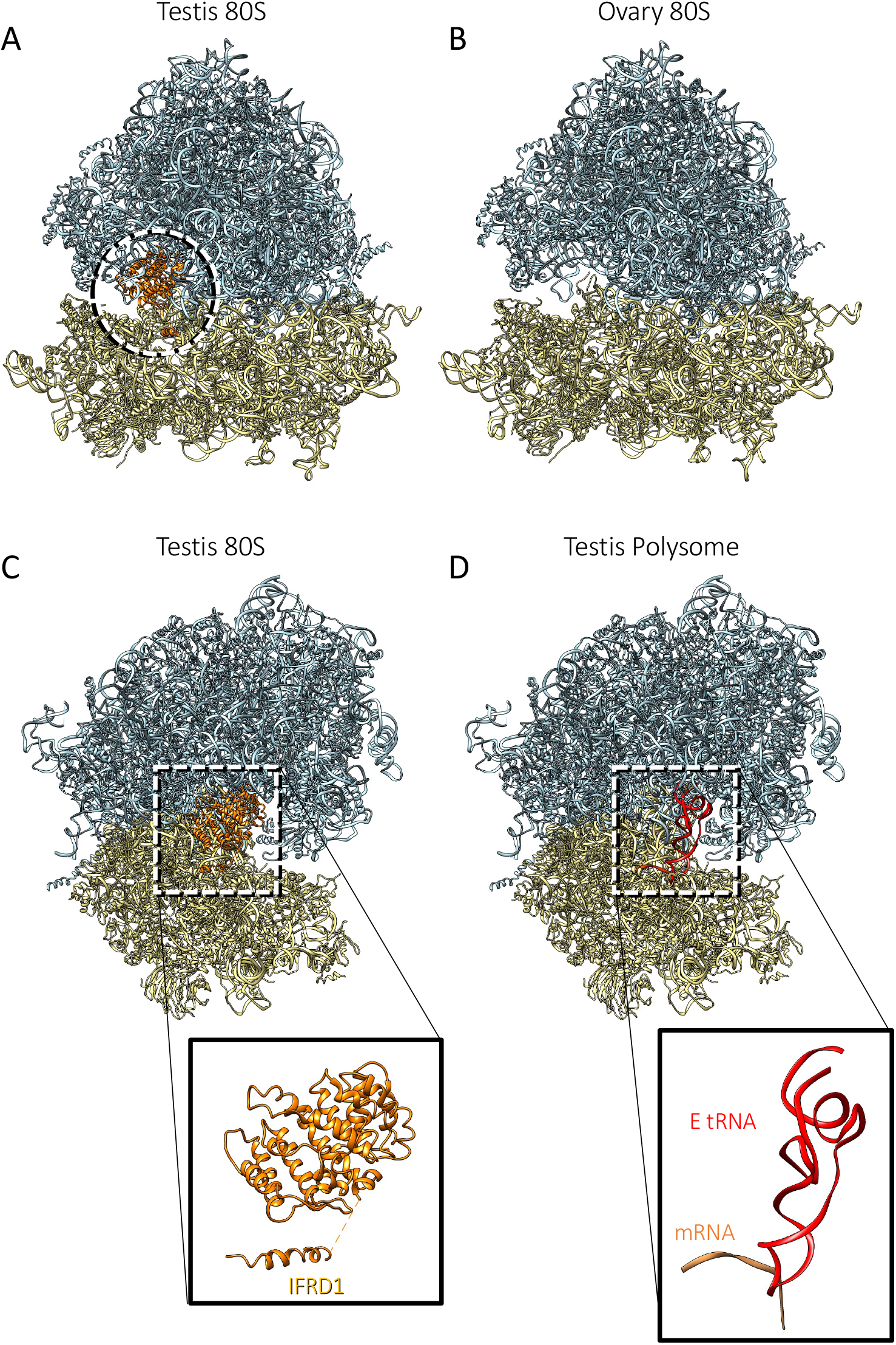
Atomic models of testis and ovary ribosomes. Atomic models of testis 80S (A and C), ovary 80S (B) and testis polysome (D). Large subunits are light blue and small subunits are yellow. IFRD1 is orange, mRNA is salmon and E tRNA is red.

Comparison of the testis and ovary 80S models revealed that the main difference between them is at the P- and E-sites (Fig 3A and B). While the ovary 80S average did not contain any densities in this region, the testis 80S average contained densities that did not correspond to a tRNA (Fig 3A, circle). As a comparison, the previously published *D. melanogaster* average contained densities for an E-tRNA and for elongation factor 2, neither of which are present in our maps [59]. By combining information from the testis 80S structure and the corresponding TMT data, we identified this density to be the interferon-related developmental regulator 1 (IFRD1) (Fig 3C), which is abundant in the testis 80S complexes (54^th^ most abundant protein in testis 80S-TMT1). Of note, rabbit IFRD2, orthologous to *D. melanogaster* IFRD1, was identified in translationally inactive rabbit ribosomes as being bound to P- and E-sites of ∼20% 80S isolated from rabbit reticulocytes [64]. Strikingly, in the reticulocytes the presence of IFRD2 is always accompanied by a tRNA in a noncanonical position (termed Z-site). In the testis 80S average no tRNA was found in this region. In mammals (rabbits and humans), IFRD2 is thought to have a role in translational regulation during differentiation [64]. Differentiation is a key process during spermatogenesis within the testis, and in this context, it is unsurprising to have found this protein in the testis 80S. *D. melanogaster* IFRD1 has considerable amino acid sequence conservation with rabbit IFRD2 (37% identity, Sup 11A and B). The presence of IFRD1 suggests that a significant proportion of the testis 80S ribosomes are not actively engaged in translation. The IFRD1 density was not present in the ovary 80S structure, suggesting far fewer ribosomes are inactive by this mechanism in the ovary. The presence of IFRD1 does not affect the paralog enrichment events because the testis specific paralog enrichment was identical between the testis 80S and testis polysome ribosomes, and polysomes are unlikely to be translational repressed. To verify this, we solved the structure of ribosomes isolated from testis polysomes (cryo-EM average resolution was 4.9 Å) (Fig 3D and Sup 10G-I). It is clear from the density map that IFRD1 is not present in either the P- or E-sites; rather there is density for the E-tRNA in these actively translating ribosomes (Fig 3C and D). In summary, we have solved the structures of testis 80S, ovary 80S and testis polysomal ribosomes, which exhibit differences in translational activity.

### Functional implications of paralog switching event in gonads

To understand the implications of the enrichment for different RP paralogs between testis and ovary 80S ribosomes we probed our two 80S cryo-EM structures. By mapping the 12 *Drosophila* paralog pairs onto our ribosome structures (all paralog pairs except RpS14a and RpS14b, as they have the same amino acid sequences), we identified three clusters in which they localise. 1) Paralogs within the small subunit, including RpS28a/b, RpS5a/b and RpS19a/b, map to the head of the 40S near the mRNA channel (Fig 4A and B). 2) Paralogs within the large subunit tend be surface-exposed. Specifically, RpL22/RpL22-like and RpL24/RpL24-like locate towards the back of the ribosome (Fig 4C-E). 3) Paralogs that are located in ribosome stalks, RpLP0 and RpL10A, potentially interacting with the mRNA during translation (Fig 4F). Of note, several small subunit paralogs are close to the mRNA channel, pointing towards possible functional differences in mRNA selectivity of the ribosome.

**Figure 4:**
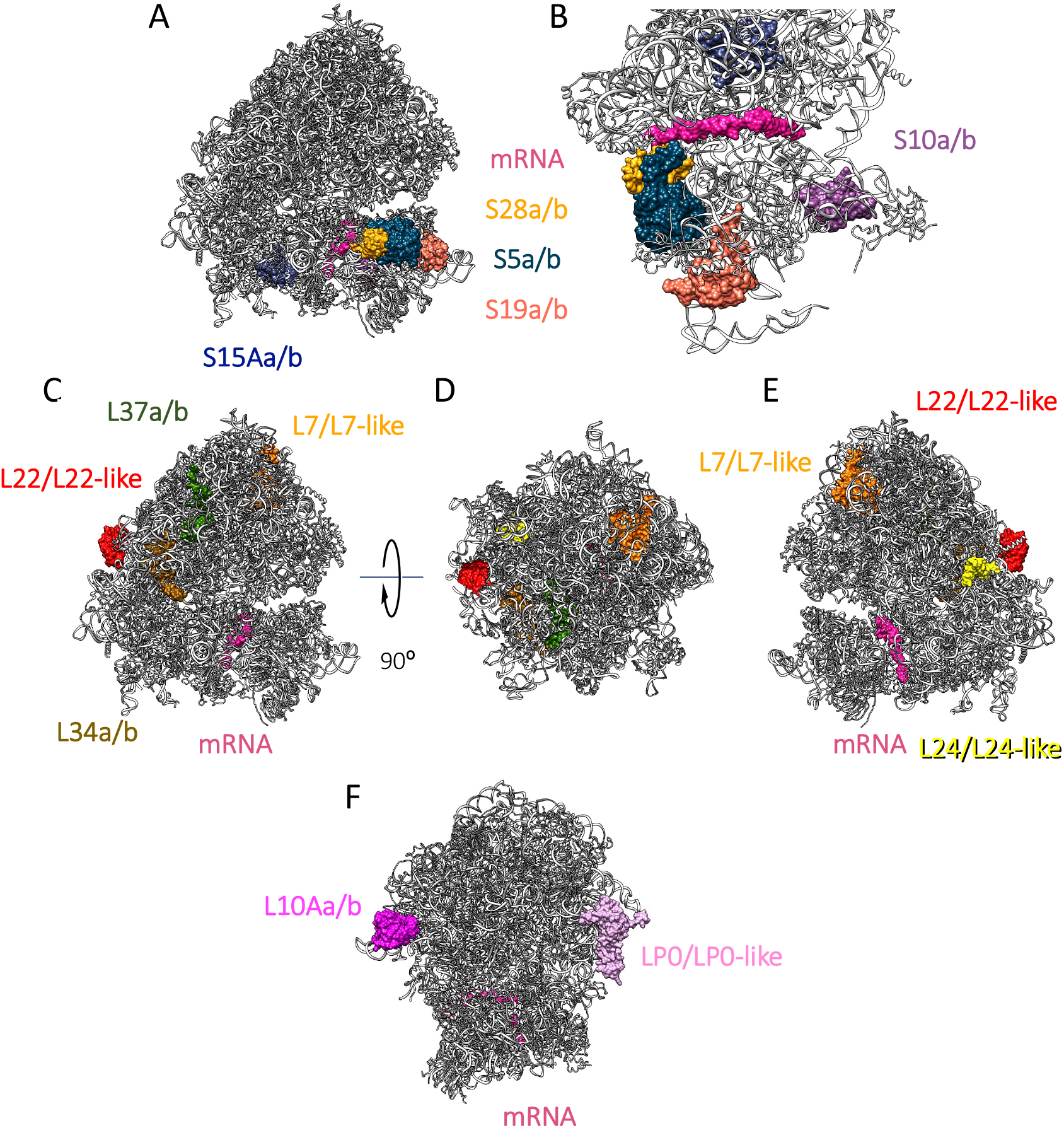
Location of *D. melanogaster* ribosomal paralogs. Ribosomal paralogs mapped to the testis 80S EM average. (A and B) Small subunit paralogs: RpS15a/b (medium blue), RpS28a/b (gold), RpS5a/b (navy blue), RpS19a/b (coral) and RpS10a/b (purple); mRNA is pink. Paralogs are shown viewed from the side into mRNA channel (A) and from the top, without the large subunit (B). (C-E) Large subunit paralogs: L37a/b (green), L22/L22-like (red), L7/L7-like (orange), L34a/b (brown) and L24/L24-like (yellow) ; mRNA is pink. Paralogs are viewed from the side into mRNA channel (C), from the top of the ribosome (D) and from the opposite side of the mRNA channel (E). (F) Paralogs that locate in ribosome stalks: L10Aa/b (dark pink) and LP0/LP0-like (light pink); mRNA is pink. Paralogs are shown viewed from the front of ribosome.

By comparing the atomic models for testis 80S and ovary 80S, switching the paralogs identified by western blot, we found small differences in the paralog positions between ovary and testis ribosomes (Table 1). Specifically, the three switched paralogs (RpL22-like and RpL37b in testis 80S; and RpS5b in ovary 80S; Fig 2D) showed the largest differences in their atomic models out of all paralogs (Fig 5A-I). Additionally, analysing the fit of both RpL22-like and RpL22 into the cryo-EM density of testis 80S ribosomes showed a better fit for RpL22-like, further pointing towards a paralog switch (Sup 12). This was also the case for L37b in testis 80S ribosomes (Sup 13). Of the paralogs we could not confirm a switch between testis 80S and ovary 80S by western blot, RpS19a, RpS10b and RpS28b showed the largest differences (Sup 14 and 15). We note that the differences of the atomic models of these paralogs between testis and ovary 80S ribosomes is above the differences for non-paralog models, that were used as control (Table 1). These differences might represent actual paralog switches or could be due to the low resolution at the head of the small subunit. For RpS5a and RpS28b the differences could also be due to their proximity to the E- and P-sites and therefore the position of IFRD1 in the testis 80S (Sup 16).

**Figure 5:**
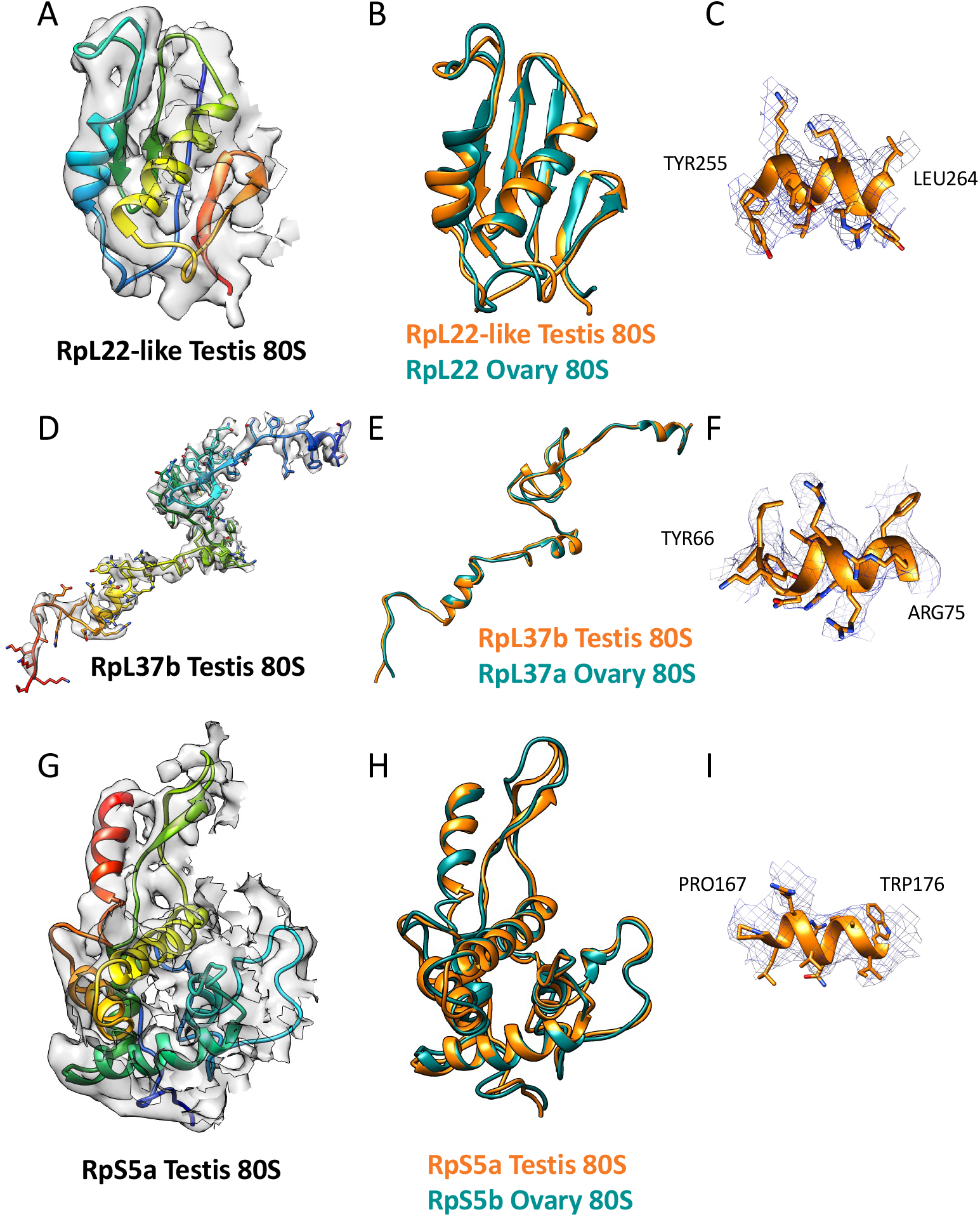
Structural implications of paralog-switching events. Switched paralogs in testis 80S vs ovary 80S are shown. (A-C) RpL22-like (testis 80S) and RpL22 (ovary 80S). (D-F) RpL37b (testis 80S) and RpL37a (ovary 80S). (G-I) RpS5a (testis 80S) and RpS5b (ovary 80S). (A, D and G) Testis 80S atomic model fitted into the EM density. Models are rainbow colored from N-terminus (blue) to C-terminus (red). (B, E and H) Comparison between the testis 80S (orange) and the ovary 80S (teal) atomic models. (C, F and I) Representative fits of the testis 80S atomic models into the EM map.

Comparing the amino acid sequences of each paralog pair it is possible to predict that they might contribute different functionality to the ribosome (Table 2 and Sup 17). RpL22 and RpL22-like are only 45% identical, even though they are very similar in length (Fig 6A, Sup 17). Unfortunately, the most different region between RpL22 and RpL22-like (i.e., the N-terminal region; Fig 6A), faces the exterior of the ribosome and is not resolved in the cryo-EM density (Fig 6A shows in bold the regions of RpL22 and RpL22-like present in the ovary 80S and testis 80S reconstructions, respectively). It is possible to imagine that given the majority of these paralogs are localised to the exterior of the ribosome, by switching one for the other might provide a difference exterior surface, with which other associated factors might bind.

**Table 2:** Summary of paralog pair attributes. For each paralog pair: amino acid length, amino acid identity, tissue enrichment or switching, relationship to human RPs and associated human diseases.

| Canonical paralog | aa length | Alternative paralog | aa length | Paralog homology | Tissue enrichment | Human ortholog(s) | Human Disease |
| --- | --- | --- | --- | --- | --- | --- | --- |
| RpL22 | 299 | RpL22-like | 312 | 45% | Testis RpL22-like | RpL22<br>RpL22L1 | Cancer/blood disease |
| RpL37a | 93 | RpL37b | 89 | 75% | Testis RpL37b | RpL37 |  |
| RpS19a | 160 | RpS19b | 159 | 66% | Testis RpL19b | RpS19 | Diamond-Blackfan anaemia. Cancer. |
| RpS10a | 163 | RpS10b | 160 | 61% | Testis RpS10a | RpS10 | Diamond-Blackfan anaemia. Aase syndrome |
| RpS28a | 64 | RpS28b | 65 | 82% | Testis RpS28a | RpS28 | Diamond-Blackfan anaemia. |
| RpS15Aa | 130 | RpS15Ab | 130 | 97% | Testis RpS15Ab | RpS15A | Diamond-Blackfan anaemia. |
| RpL24 | 155 | RpL24-like | 191 | 24% | Ovary 80S RpL24-like | RpL24 | Tropoblast development |
| RpL7 | 252 | RpL7-like | 257 | 28% | Ovary 80S RpL7-like | RpL7 | Cancer |
| RpLP0 | 317 | RpLP0-like | 256 | 23% | Ovary 80S RpLP0-like | RPLP0 |  |
| RpS5a | 228 | RpS5b | 230 | 76% | Ovary>Embryo><br>Testis RpS5b | RpS5 | Diamond-Blackfan anaemia. Cancer. |

**Figure 6:**
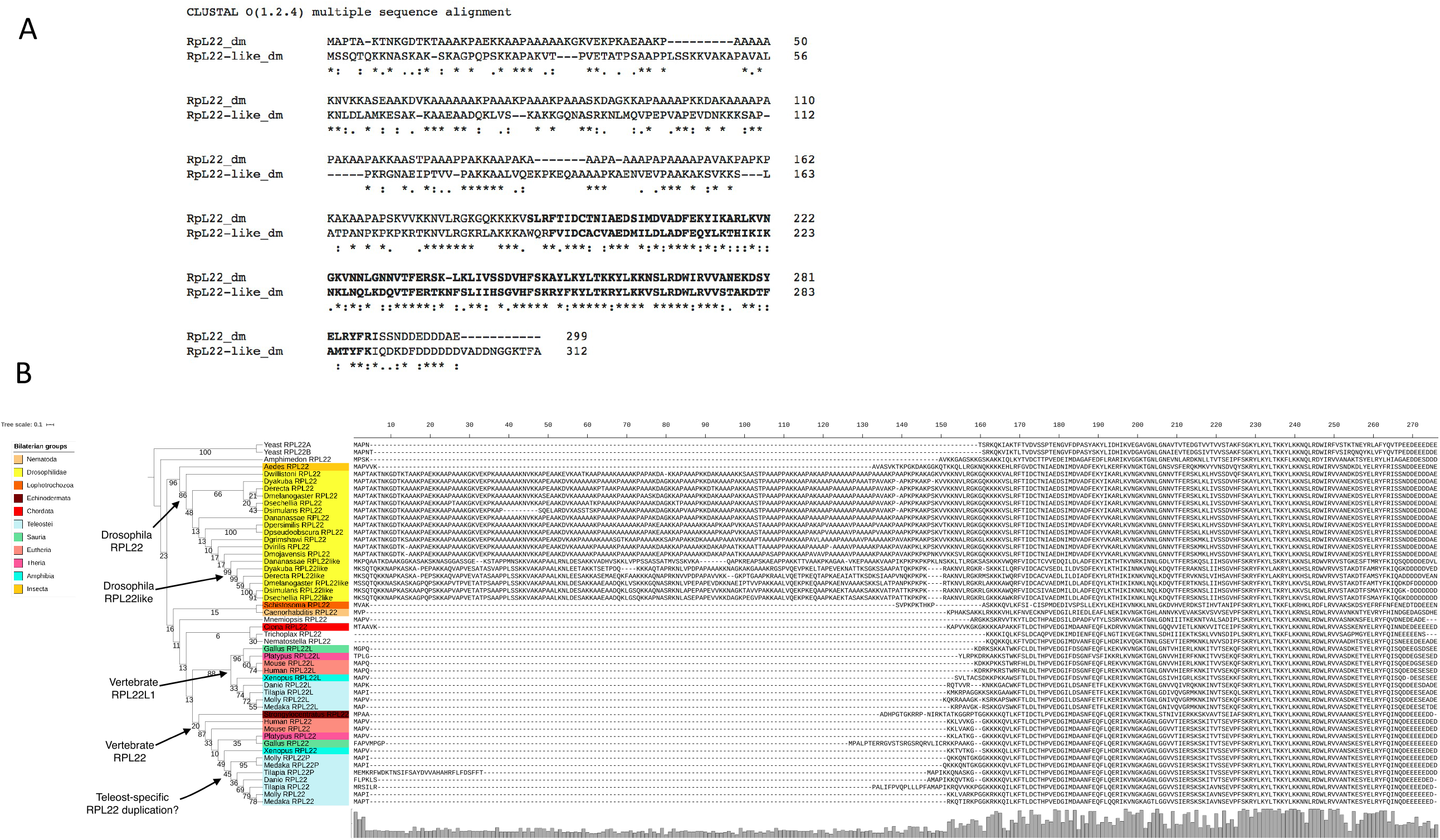
Evolution of RpL22 paralogs by independent duplication events. (A) Alignment of RpL22 and RpL22-like amino acid sequences for the two *D. melanogaster* proteins. In bold are amino acids included in the atomic models. (B) Phylogenetic tree for RpL22 and RpL22-like paralogs across a range of animal genomes. The three duplication events detected are indicated.

### Evolution of RpL22 paralogs by independent duplication events

To probe how widespread paralog switching events might be to facilitate ribosome specialisation we determined the level of conservation of RpL22 and RpL22-like in other animal genomes. Orthologs of RpL22 were identified across a range of animals including Drosophilid*s*. We determined that the paralogous pair RpL22 and RpL22-like present in *D. melanogaster* evolved by 3 independent duplication events across the animal clade (Fig 6B). A duplication event unique to the Drosophila clade produced the paralogous pair RpL22 and RpL22-like that are identifiable in 6 out of the 12 Drosophila species sampled. The additional 2 duplication events are present in the vertebrate clade and may be the result of whole genome duplication rather than individual gene duplication events. The first of these vertebrate duplications produced the paralog pair RpL22 and RpL22L we observe in humans for example. The second vertebrate *RPL22* duplication specific event occurred amongst teleost fishes and the most parsimonious explanation of pattern of distribution of duplicate copies would suggest subsequent loss in some lineages (Fig 6B). Thus, *RPL22* has undergone multiple independent duplication events, generating a complex array of paralogous pairs.

## DISCUSSION

We have characterised the heterogeneity of 80 S ribosome composition across ribosomes purified from 4 *in vivo* tissues The main source of heterogeneity we discovered were paralog-enriching events in the gonads. We have identified 6 testis-enriched paralogs (RpL22-like, RpL37b, RpS19b, RpS10a, Rp28a, RpS15Ab) and 4 ovary-enriched paralogs (RpL24-like, RpL7-like, RpL0-like and RpS5b). In addition to being ovary-enriched RpS5b is also present in embryo and testis but to a lesser extent. We were able to validate testis-enrichment for RpL22-like, RpL37b and ovary-enrichment for RpS5b by western blots. These westerns also suggest that these alternative paralogs are more abundant than the canonical ones, indicating that paralog-switching is occurring, not simply paralog-enrichment. Furthermore, the enrichment of RpS19b and RpS5b in the testis is to such an extent that they are at about equal levels to their respective canonical pairs, i.e ∼50% of ribosomes likely contain RpS19b and RpS5b in the testis. There are very few differences between the composition of 80S and polysome ribosomes across all tissues. Paralog incorporation is not simply the consequence of transcriptional regulation of these paralogous genes. Rather there is modulation at the level of the translation of some of these proteins and/or incorporation into the ribosome. We can be sure that the difference we have detected are within ribosomes, rather than other large RNPs, as EDTA treatment disrupts our complexes.

We have solved the cryo-EM structures of three different ribosome complexes purified from complex *in vivo* tissues; 80S ribosomes from the testis (3.5 Å), 80S ribosomes from the ovary (3.0 Å) and polysomal ribosomes from the testis (4.9 Å), improving the resolution from the only other previous ribosome from *D. melanogaster* [59]. One key difference was that the testis 80S structure contains IFRD1, which is a homolog of the human IFRD2. Its presence indicates there is functional homology between the two proteins in inhibiting mRNA translation through the ribosome, during differentiation. In mammals IFRD2 was seen in differentiating reticulocytes [64], whilst in our work we found IFRD1 in the testis 80S (but not in the ovary 80S), where it could be involved in regulation of translation during the differentiation of spermatozoa, which is central to the function of the testis.

*D. melanogaster* paralogs are localised in three clusters; a) the head of the 40S near the mRNA channel, b) the surface-exposed back of the large subunit and c) ribosome stalks, potentially interacting with the mRNA during translation. The position of these three clusters provides potential explanations of how specialisation might be achieved, mechanistically. Differences in amino acid sequence and precise position of the testis and ovary switched paralogs (Fig 5) could potentially affect the interaction of the mRNA and the ribosome, specifically during initiation when 40S ribosomes are recruited to the 5’ end of mRNAs. The back of the 60S would provide an ideal site for additional protein factors to differentially bind to ribosomes containing these RPs, with the potential to regulate ribosome activity. This is particularly true for the RpL22 and RpL22-like paralog pair, which has the lowest sequence identity between each other, 45%. The termini of these proteins are likely to be dynamic given the lack of density for them in our EM maps. Our phylogenomic analysis suggests that the modulation of this part of the exterior ribosome surface is in common across many organisms, and that the generation of paralogs has occurred independently three times for RpL22.

Therefore, this provides a potential mechanism for ribosome regulation across many eukaryotes. Although paralogs are not conserved across a range of organisms, and many are limited to Drosophilids, there are many organisms with many RP paralog pairs, including human (19 pairs) and *Arabidopsis* (80 pairs). Therefore, these potential mechanisms of ribosome regulation could be conserved, if not the precise details.

mRNA translational regulation is important in the testis and ovary. For example, many testis-specific translation components exist to enable tight regulation such as eIF4-3 [29]. The result we find here, that the gonads are important sites of ribosome heterogeneity, suggests that RP paralog enriching, and potentially paralog switching, might also play a part in this regulation via ribosome specialisation. The paralog enrichment events in the testis we have identified involve pairs where the ‘canonical’ paralog in the pair is located on the X chromosome. One hypothesis for the expression of the ‘alternative’ paralog in testis enables gene dosage to be maintained during meiotic sex chromosome inactivation, which occurs in males [65]. However, additional functions could have evolved as well as this.

The importance of the paralog-switching event between RpS5a and Rp5b has recently been functionally characterised in the *Drosophila* ovary [34]. Females without RpS5b produce ovaries with developmental and fertility defects, whilst those without RpS5a have no defects. RpS5b specifically binds to mRNAs encoding proteins with functions enriched for mitochondrial and metabolic GO terms in the ovary, suggesting ovary RpS5b containing ribosomes translate this specific pool of mRNAs [34]. It will be interesting to see how widespread this finding is for RpS5b in other tissues, since this is a frequently enriched paralog; it is enriched in embryo and testis to a lesser extent. It has been known for some time that mutations in RpS5a produce a *Minute* phenotype (including infertility), so it seems likely that these two paralogs each have biologically important roles in the fly. RpS5a and RpS5b have also been seen to exhibit tissue-specific expression in *A. thaliana*, in a developmentally regulated manner [15]. atRpS5a was suggested to be more important than atRpS5b during differentiation, because of its expression pattern, but the regulation mechanism remains elusive in *A. thaliana*.

The function of the RpL22 and RpL22-like paralog pair in the *Drosophila* testis has been previously explored and it has been suggested that the two proteins are not functionally redundant in development or spermatogenesis [66, 67]. Further work is needed to directly link effects on ribosome composition and mRNA translational output, as the two paralogs interact with different pools of mRNA in the testis [67].

Interestingly, we found little differences between 80S and polysomal ribosome composition, apart from an enrichment of RpL24-like in 80S ribosomes in the testis and head. RpL7-like was also enriched in testis 80S and RpL38 in the ovary polysome. Eukaryotic orthologs of RpL24-like are thought to have a role in the formation and processing pre-60S complexes, with RpL24 replacing RpL24-like at the very end of processing [68]. Given that we saw enrichment of RpL24-like in 80S compared to polysomes in the testis and the head, this suggests that a proportion of these 80S complexes could represent the final stage of testing 80S competency. It is not clear why this would be the case in only these tissues. RpL24-like is present in other insects and some non-insect arthropods (FlyBase). A paralog switching event between RpL24 and RpL24-like could be important in translation initiation or indeed provide a platform for additional proteins to bind to the ribosome, given RpL24/RpL24-like is located close to RpL22/RpL22-like.

Several of the RPs that have gonad-specific paralog pairs (including RpS19, RpS5, RpS10, RpS28 and RpL22 [69, 70]) have been linked with human diseases, specifically Diamond-Blackfan anemia and cancer (Table 2). Thus, it will be important to uncover if they contribute to mRNA translation regulation and work *in vivo* using *Drosophila* could help understand how they contribute to the translation of specific mRNAs.

One of the few canonical RPs we found to be differentially incorporated was RpS11 in the head 80S ribosomes. In humans RpS11 phosphorylation has been found to be linked to Parkinson’s disease [71] and higher levels of RpS11 correlate with poorer prognosis in glioblastoma patients [72]. Therefore, understanding RpS11 levels in *Drosophila* head could provide a mechanism of future exploration for dissecting the molecular mechanisms by which RP mutations result in human disease.

Altogether our data reveal ribosome heterogeneity occurs in a tissue specific manner through differential incorporation of ribosomal paralog proteins. We further show paralog switching events in the gonads and our structural analysis has provided insights into how these switches might regulate translation mechanistically. Additionally, our evolutionary data suggest heterogeneity may represent a conserved mechanism of translation regulation across eukaryotes.

## ACKNOWLEDGEMENTS

We thank the Astbury Biostructure Laboratory (ABSL) Facility Staff for assisting with cryo-EM data collection. Electron microscopy image processing was partially undertaken on ARC3, part of the High Performance Computing (HPC) facilities at the University of Leeds. We also thank Laura Wilkinson Hewitt and Brian Jackson of University of Leeds Protein Production Facility (PPF) for RpL22 and RpL22-like antibody purification. CGPM and MJOC would like to thank The University of Nottingham for HPC facilities and the School of Life Sciences for research support. Mass spectrometry was performed by Bristol University Proteomics Facility. RpS5a and RpS5b antibodies were kindly gifted by Jian Kong and Paul Lasko [34]. We would also like to thank Pavel Baranov and Gary Loughran for insightful discussions.

---

**Sup 1:**
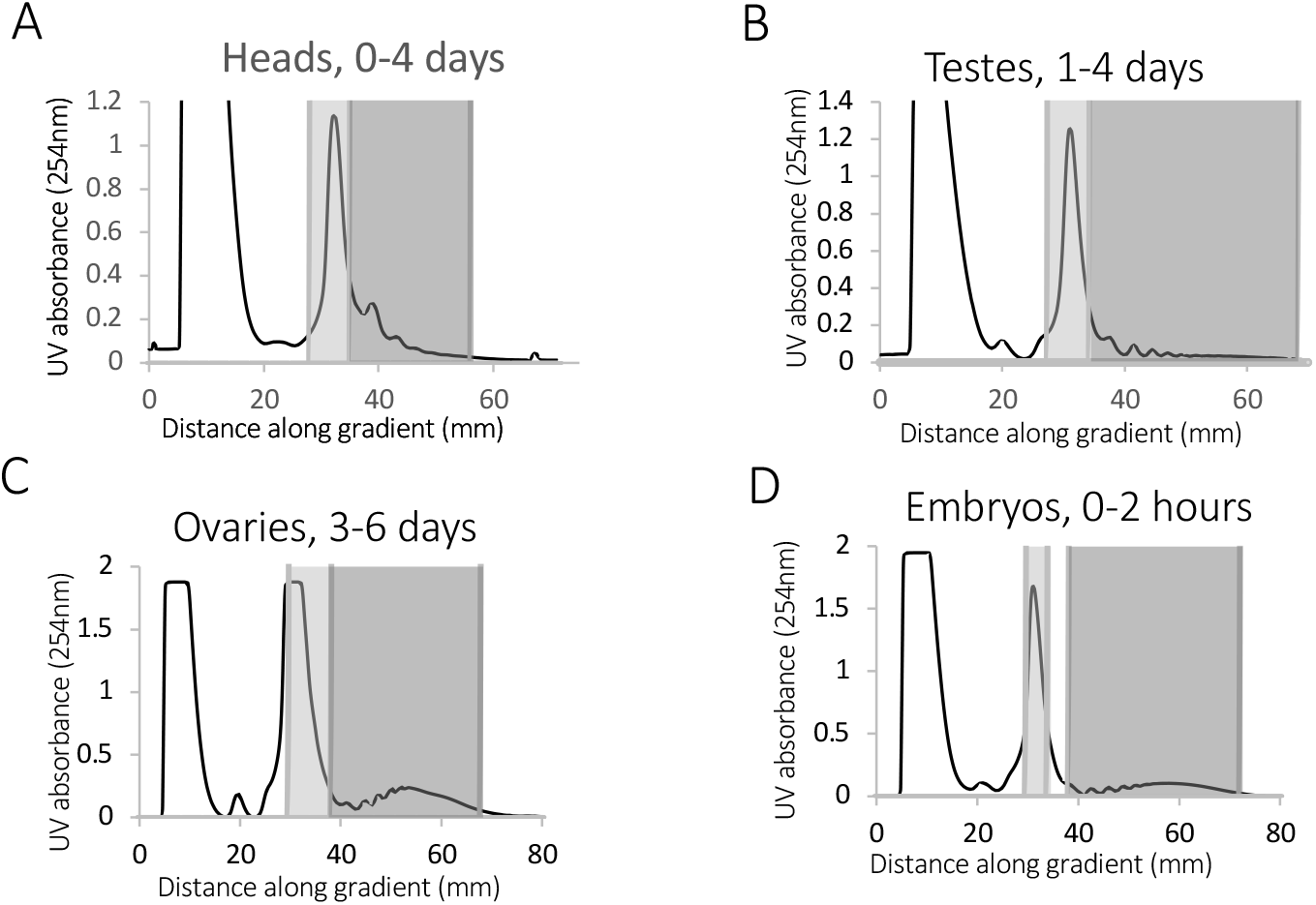
Determined ribosomal composition in tissues and during development. 254 nm UV plots across sucrose gradients with 80S and polysomal complexes isolated from (A) 50:50 mixture of female:male 0-3 day old heads. (B) ∼500 pairs of 1-4 day old adult testes, (C) ∼500 pairs of 3-6 day old adult ovaries, (D) 0-2 hour embryos. Light grey shading indicates fractions used for 80S and dark grey for polysomes.

**Sup 2:**
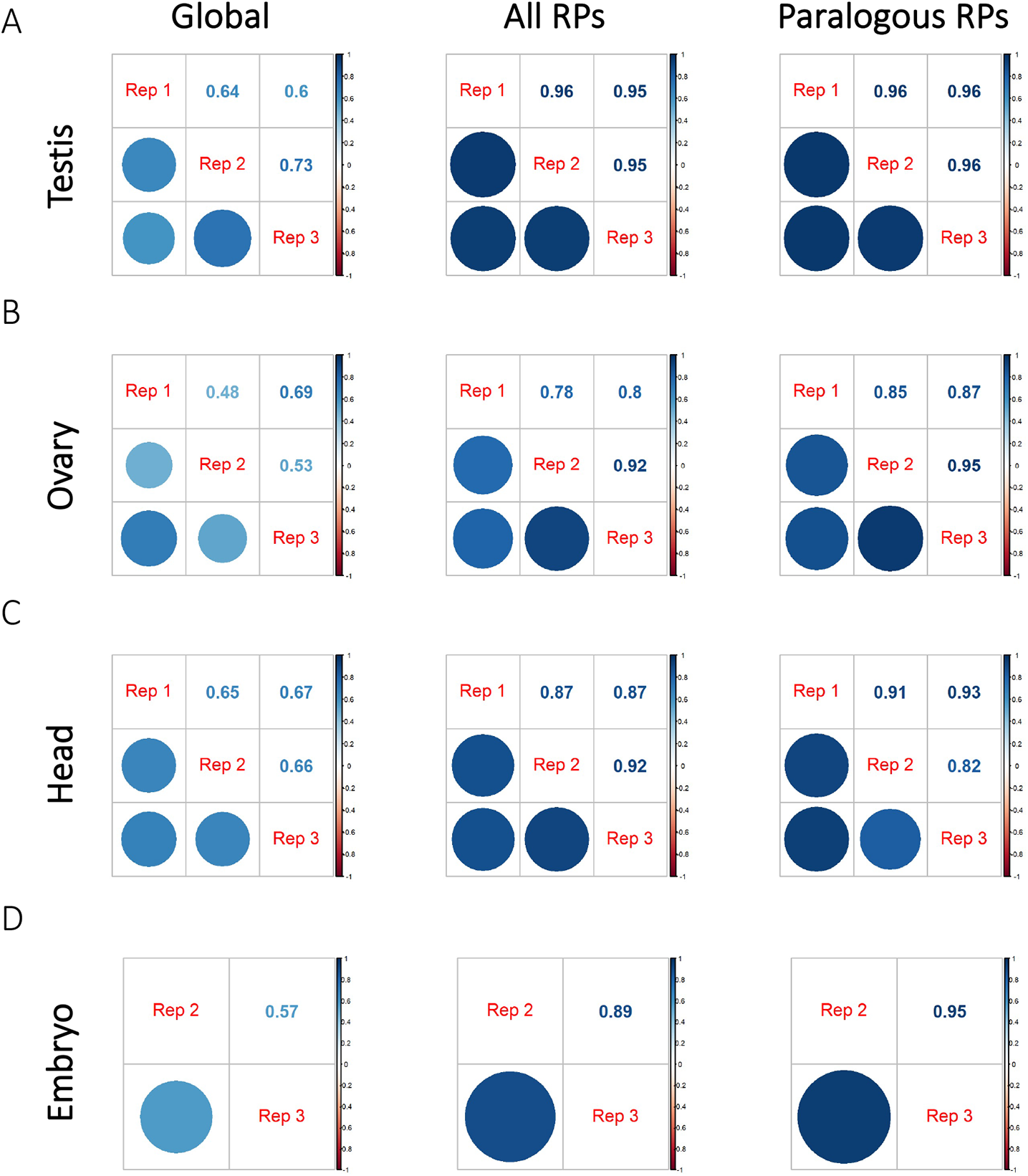
Reproducibility of TMT mass spectrometry experiments. Correlation matrices showing Pearson correlation coefficients when comparing replicates for each tissue. All *Drosophila melanogaster* proteins (global), RPs and paralogous RPs are compared from 80S monosome fractions for (A) testis, (B) ovary, (C) heads and (D) embryo tissue. Size and colour of circles correspond to correlation coefficients for all replicates, which are also stated as numbers. All comparisons were statistically significant (p-value <0.05).

**Sup 3 and 4.**
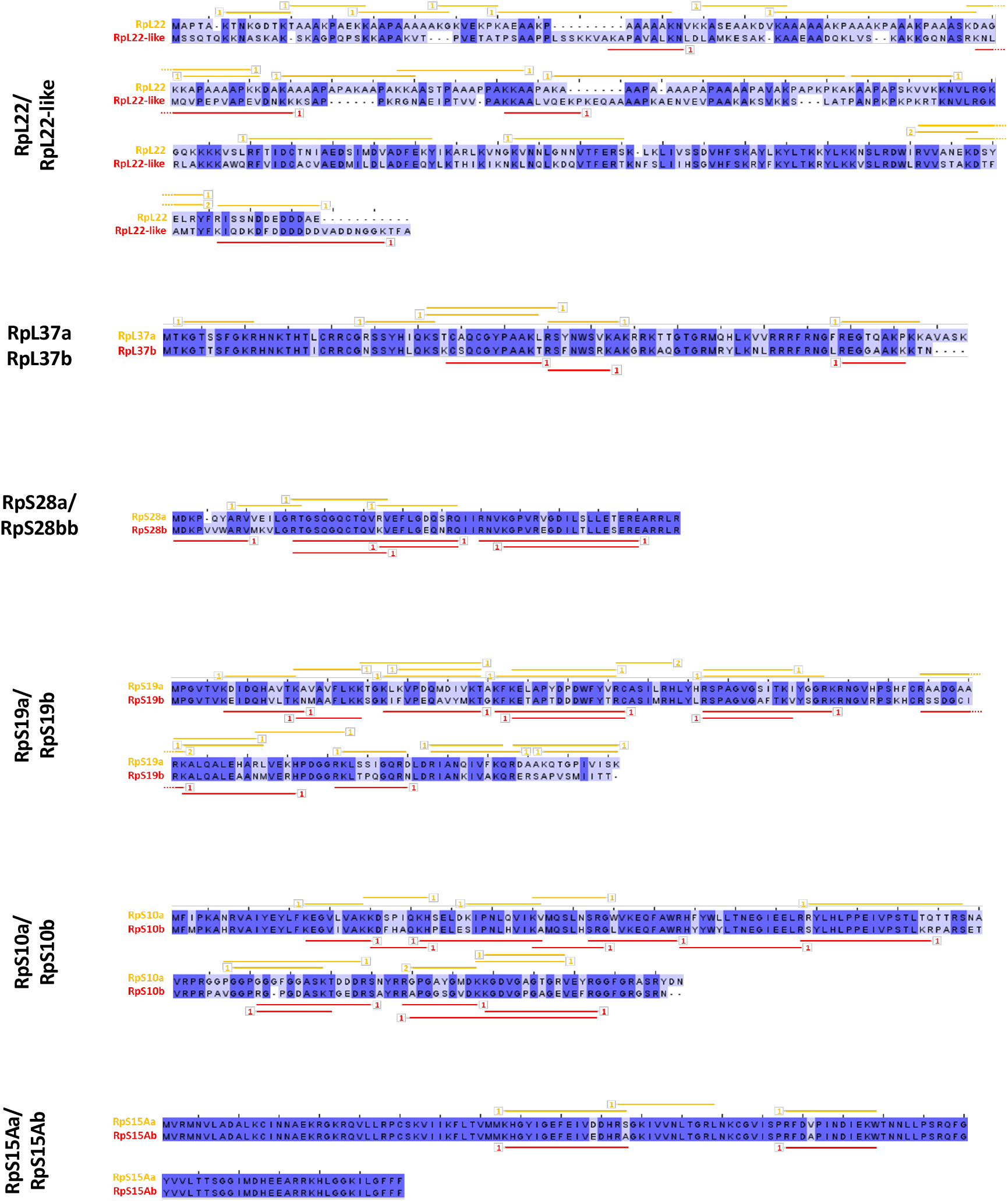
Unique peptide mapping. Clustal-omega alignment of paralog pairs with unique peptides mapped that was detected during TMT experiments. The confidence of peptide identification is given for each unique peptide: 1 denotes high confidence (<1% false discovery rate, FDR), 2 denotes medium confidence (<5% FDR).

**Sup 3 and 4.**
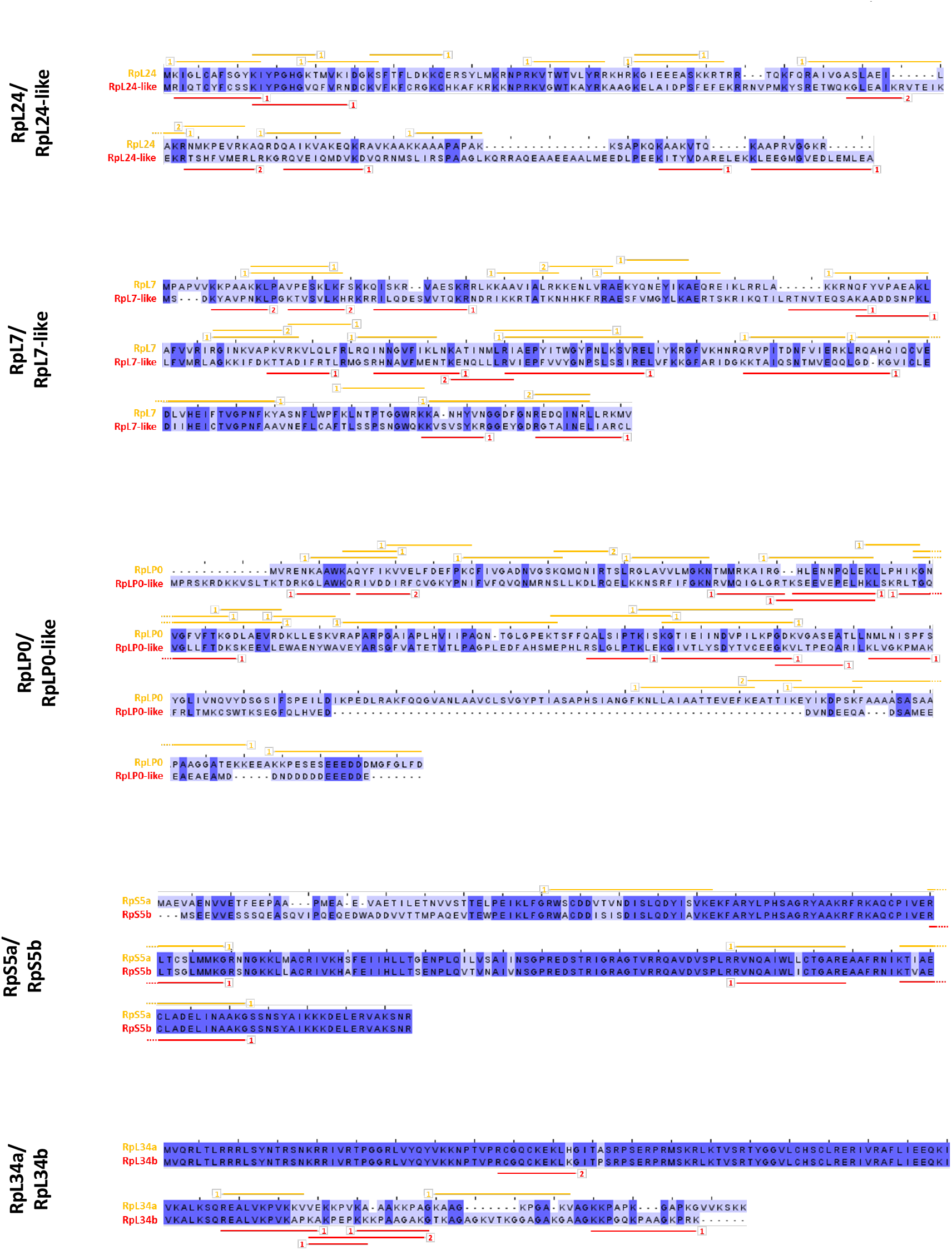

**Sup 5:**
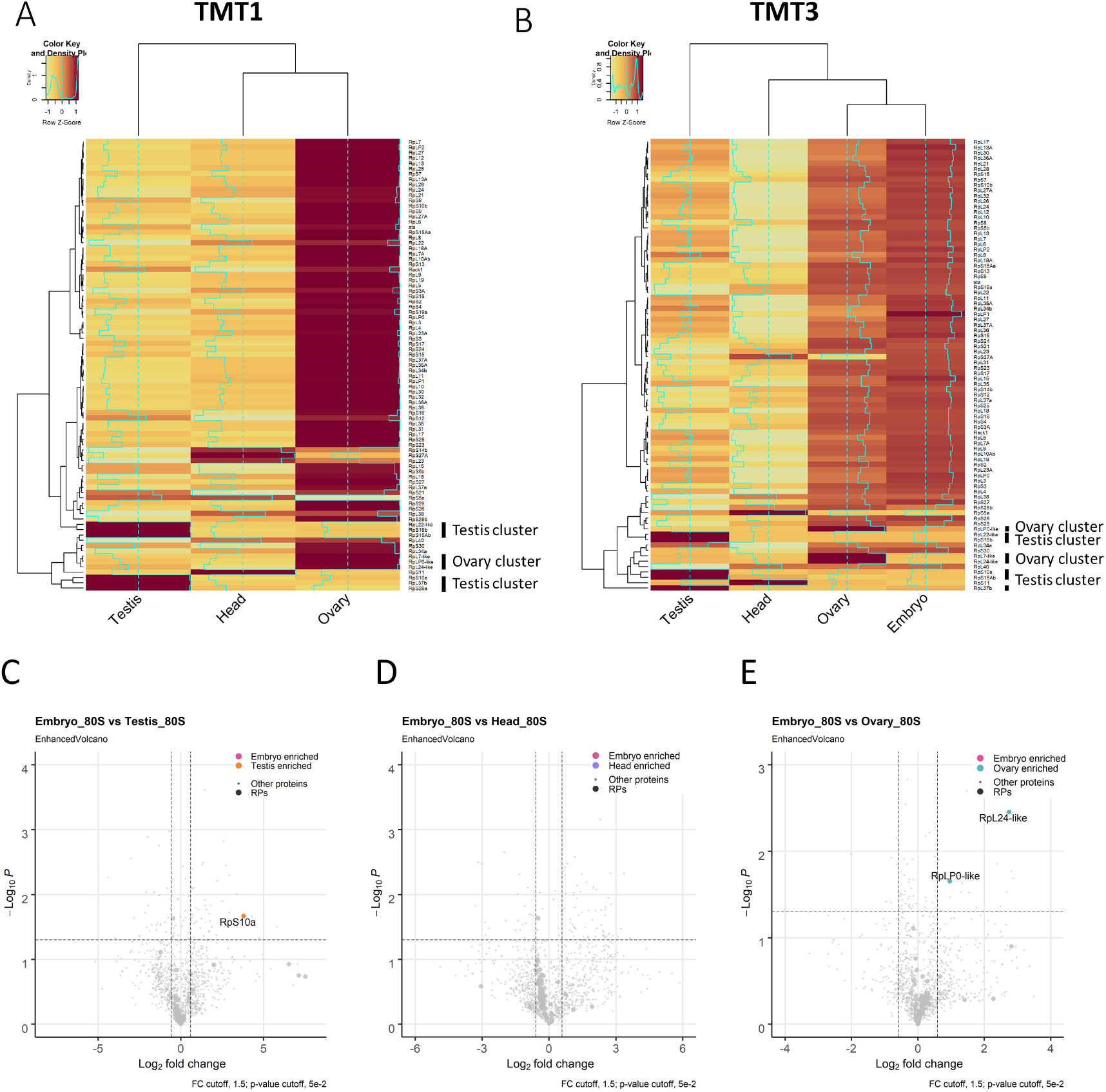
Gonad ribosome heterogeneity through paralog enrichment and paralog-switching. Hierarchical clustering of log_2_ normalised abundances from (A) TMT replicate 1 and (B) TMT replicate 3, clustered according to row. (C-E) Volcano plots highlighting little change detected by TMT-MS in the enrichment of RPs when comparing 80S from embryo to (C) testis, (D) head or (E) ovary tissue.

**Sup 6:**
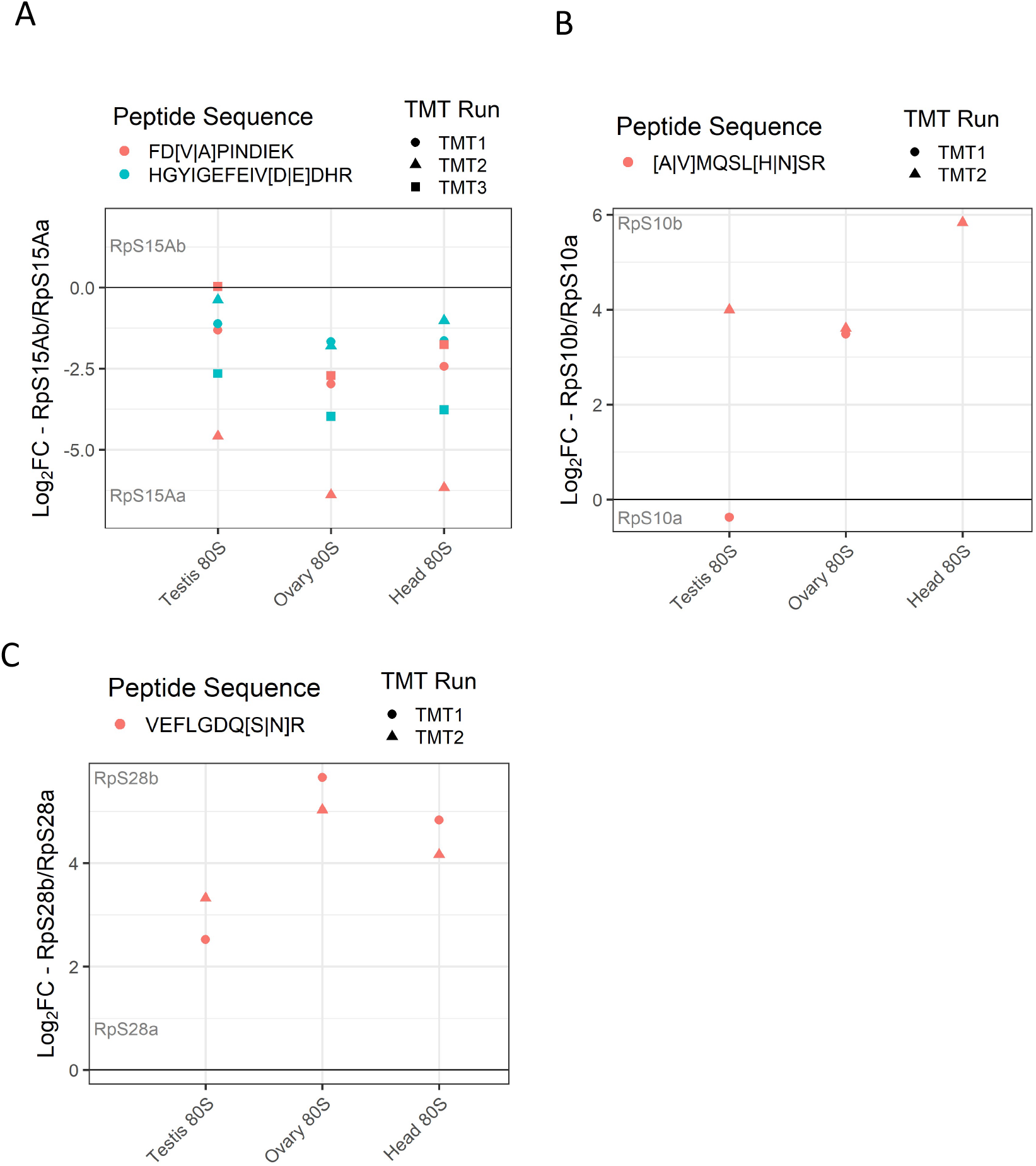
Quantification of relative paralog levels from peptide analysis. Comparison of highly similar unique peptides for (A) RpS15Aa/b, (B) RpS10a/b and (C) RpS28a/b in testis, ovary, and head tissues. Peptides compared for each paralog pair are of the same size and have 1-2 amino acid changes (differences shown in the key within square brackets). Comparisons are shown as log_2_ fold-change differences for each paralog pair.

**Sup 7:**
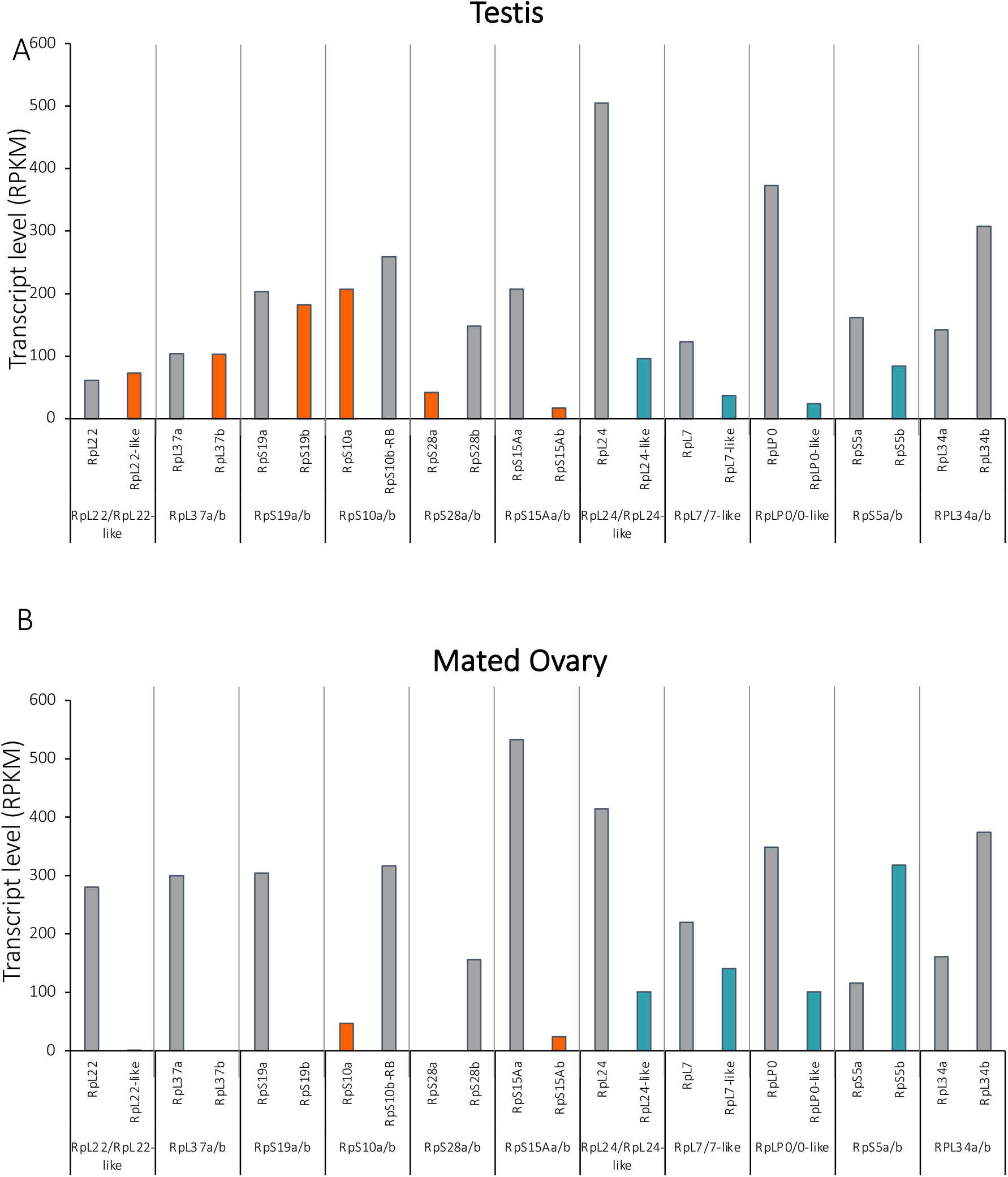
Relative levels of RP paralog pair mRNA according to RNA-Seq. mRNA levels of RP paralogs in (A) testis and (B) mated ovary tissue. RNA-Seq data was extracted from ModMine (intermine.modencode.org)(Lyne, Smith et al. 2007) with data from modENCODE project (Graveley, Brooks et al. 2011, Brown, Boley et al. 2014). Values are RPKMs. Testis enriched paralogs (orange), ovary enriched paralogs (turquoise).

**Sup 8:**
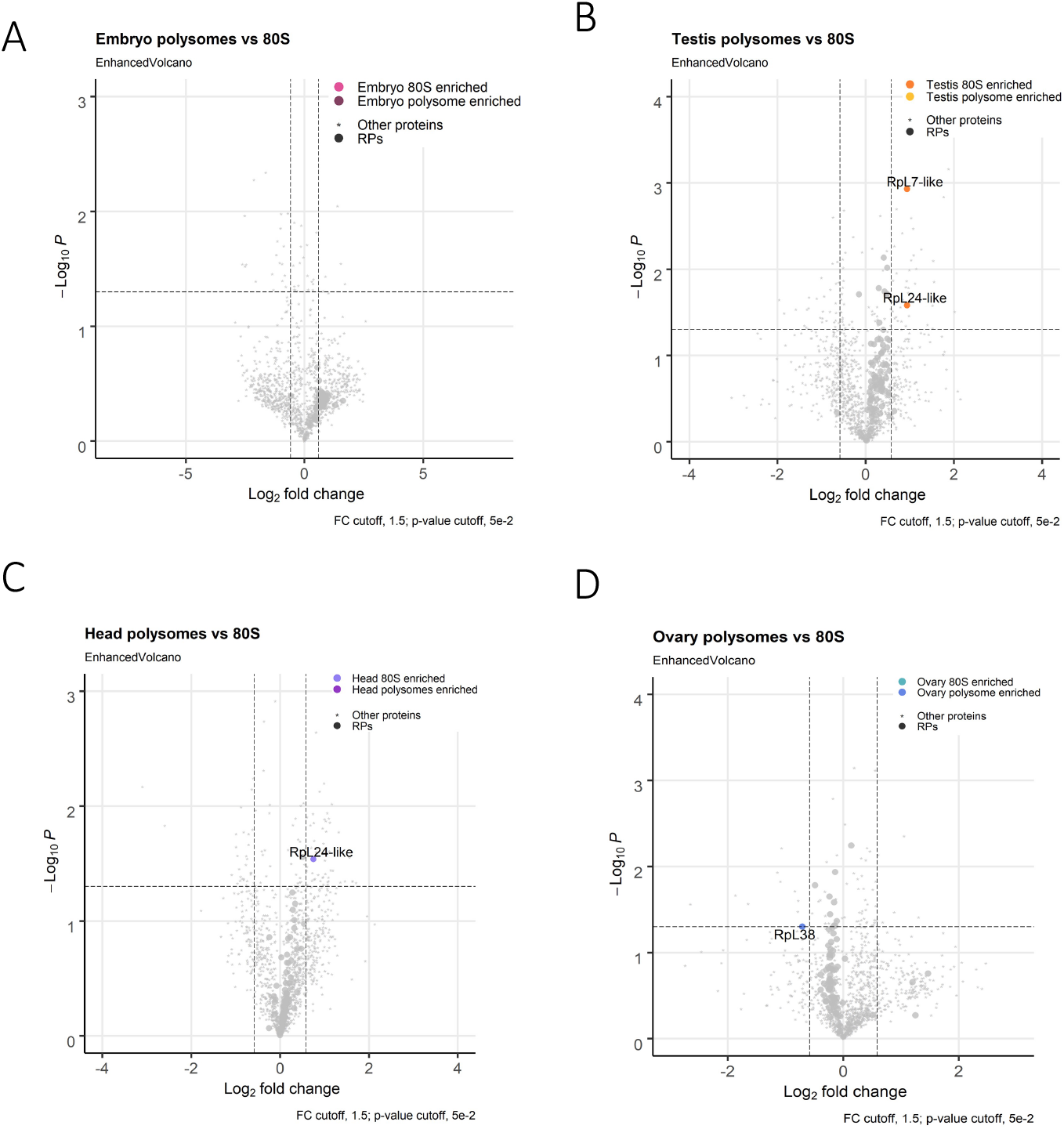
Little difference between composition of 80S and polysome ribosomes. Volcano plots comparing proteins detected in the 80S and polysome fractions for (A) testis, (B) head, (C) ovary and (D) embryo tissue. Log_2_ fold-change cut off is 1.5 and p-value <0.05. Enriched RPs are labelled.

**Sup 9:**
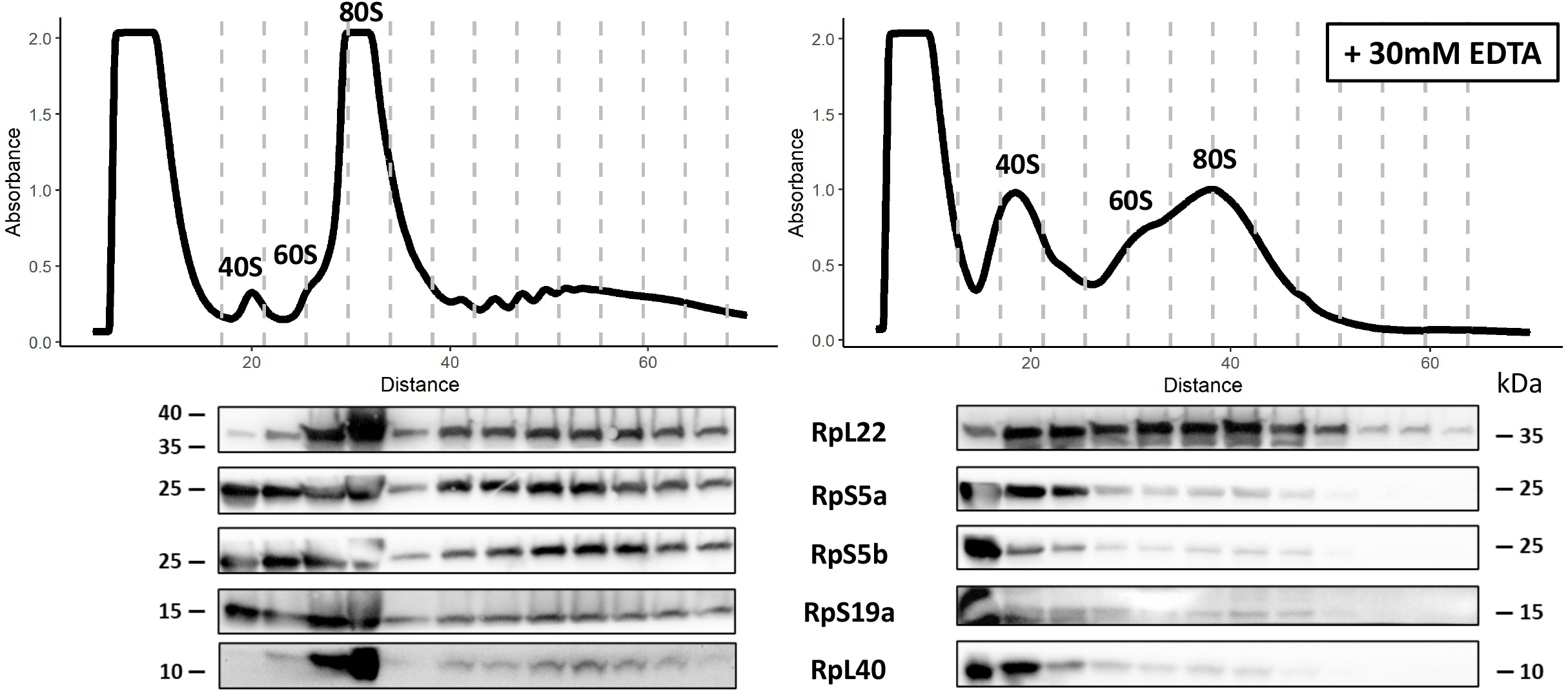
EDTA treatment indicates that enriched paralogs are part of ribosomes. 254 nm UV absorbance trace across 18-60% sucrose gradients showing isolated 80S and polysomal complexes from 150 pairs of 3-6 day old adult ovaries (A) without EDTA and (B) with 30 mM EDTA. Fractions were collected (grey lines) and subjected to western blot analysis using paralog specific antibodies. Distance refers to distance (in mm) down sucrose gradient, 0 being top of gradient (18%).

**Sup 10:**
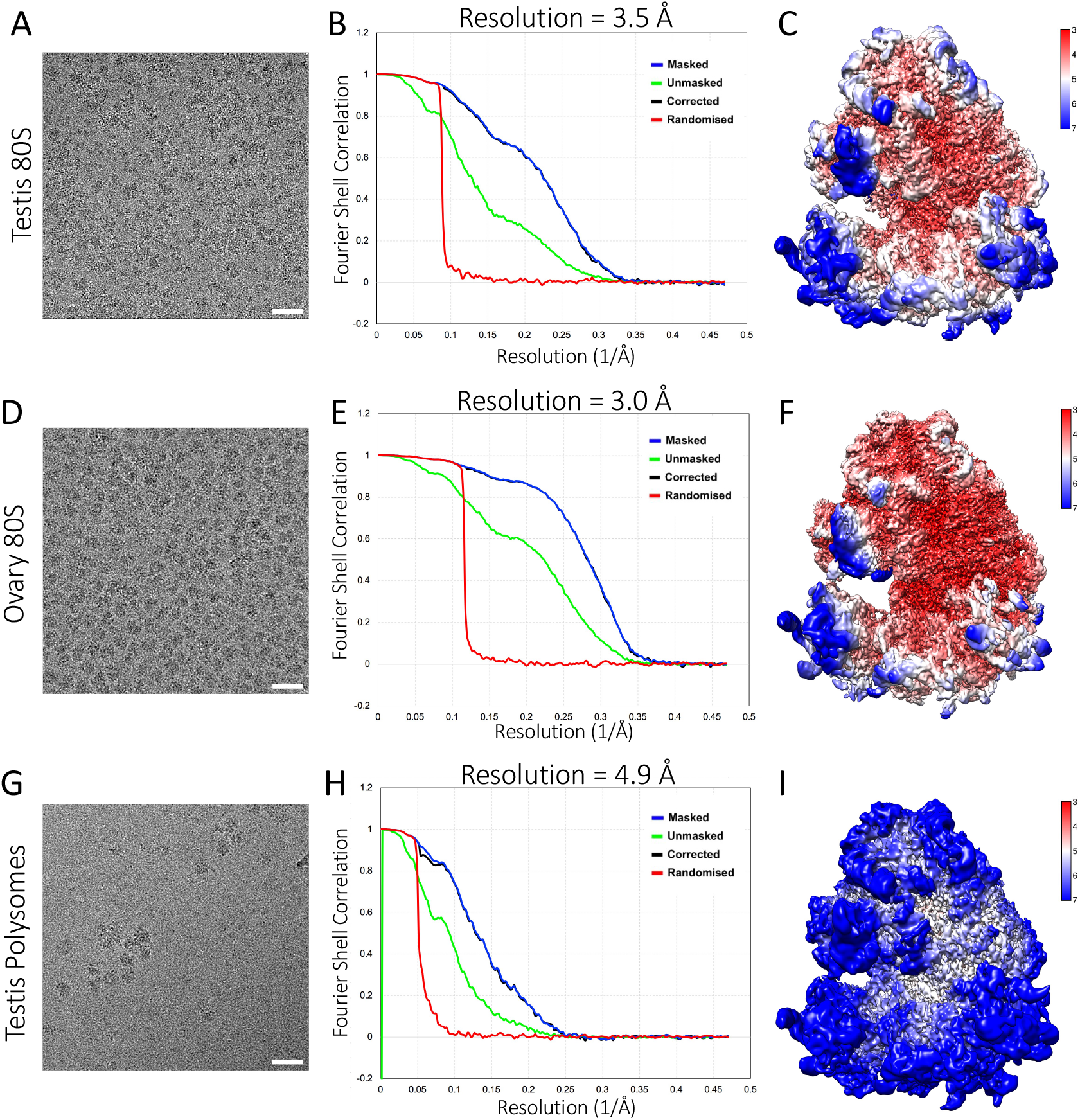
Cryo-electron microscopy of testis and ovary ribosomes. Cryo-electron micrographs (A, D and G), FSC curves (B, E and H) and local resolution coloured maps (C, F and I) for the cryo-EM averages of testis 80S (A-C), ovary 80S (D-F) and testis polysomes (G-I). Scale bars for A, D & G are 50 nm.

**Sup 11:**
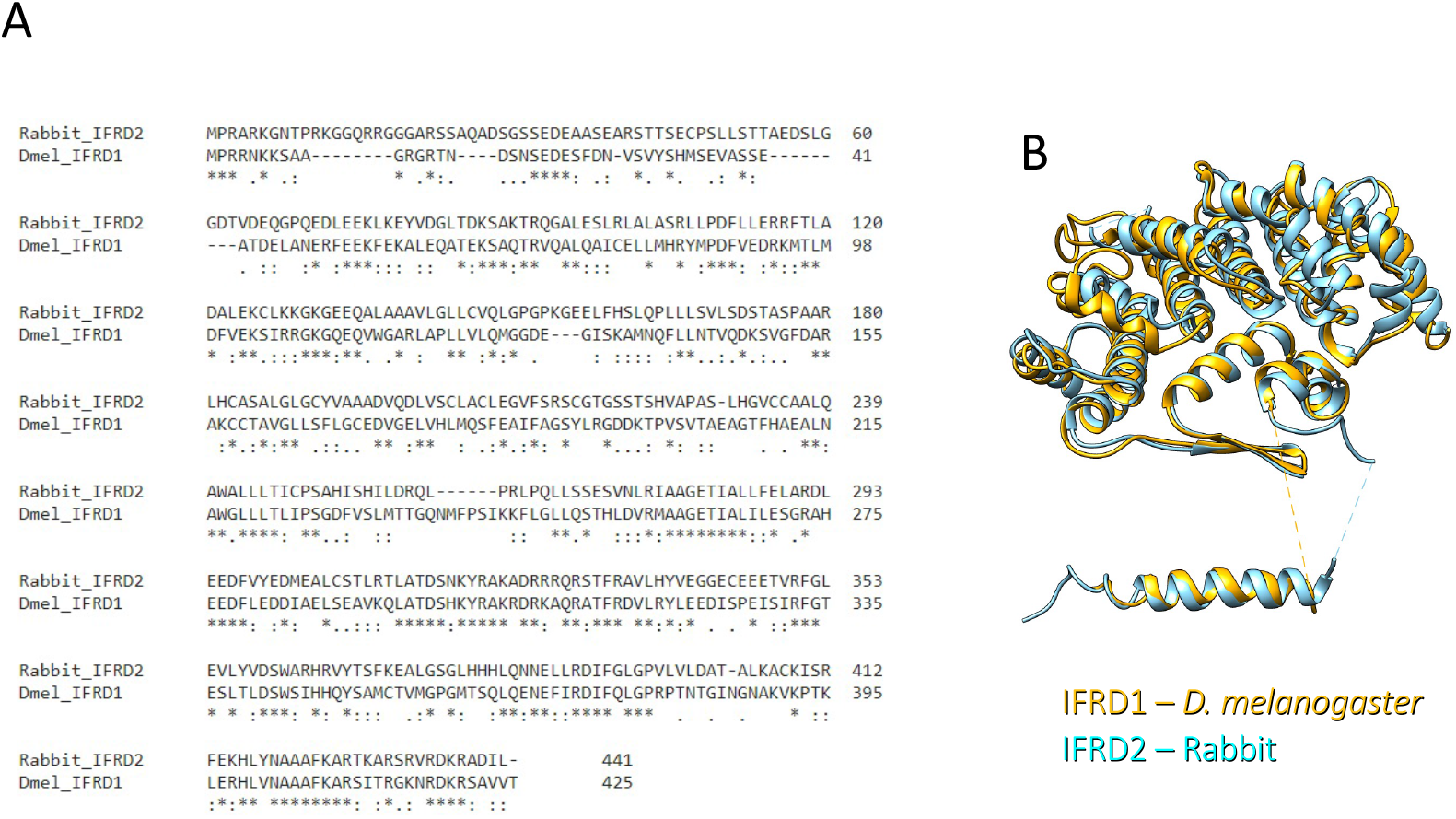
Comparison between rabbit IFRD2 and *D. melanogaster* IFRD1. (A) Clustal-omega alignment of rabbit IFRD2, and *D. melanogaster* IFRD1 protein sequence. (B) Comparison of the atomic models of rabbit IFRD2 and *D. melanogaster* IFRD1.

**Sup 12:**
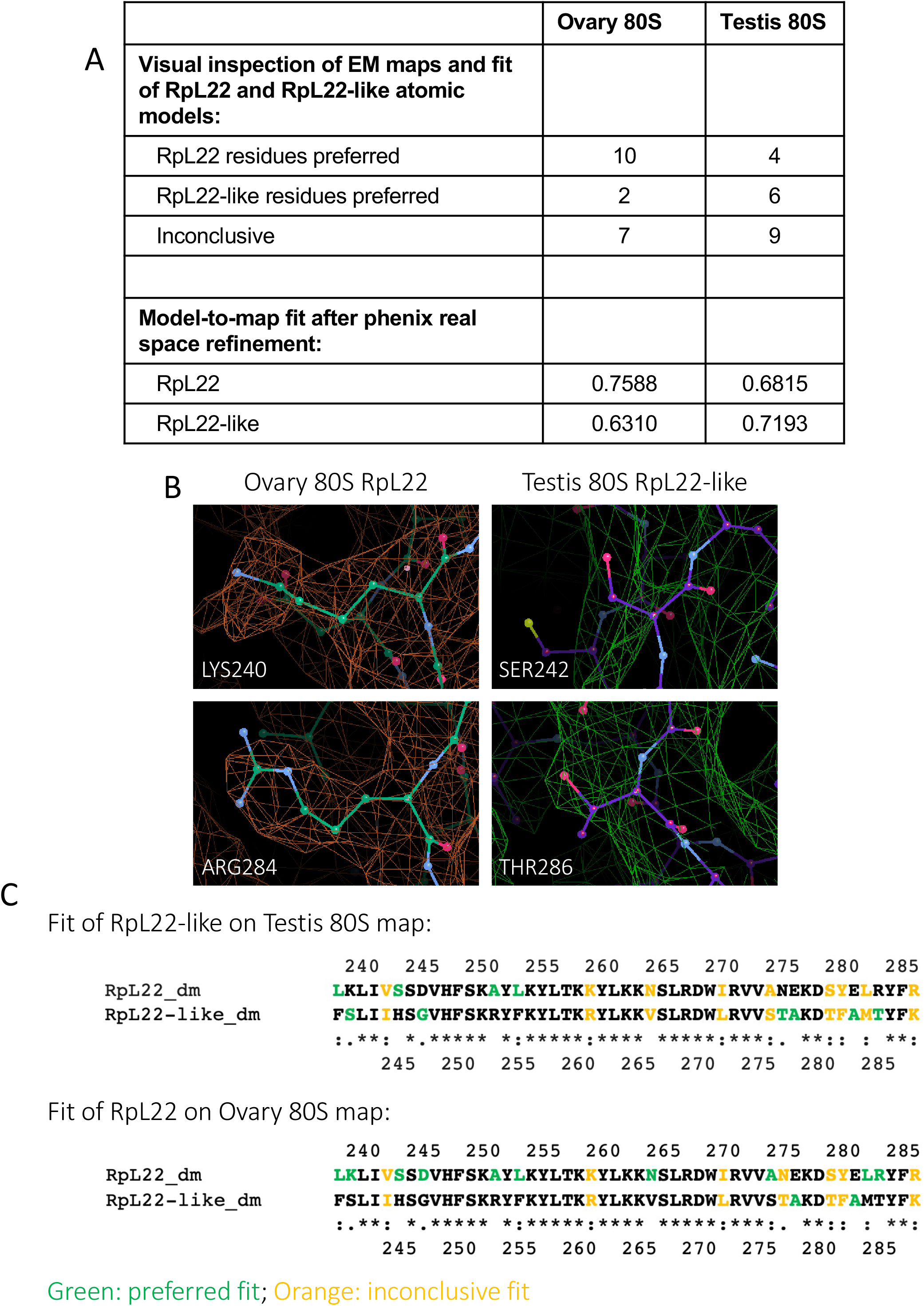
In-depth analysis of density of RpL22/RpL22-like in ovary and testis 80S EM maps. (A) Summary of the fit of residues for RpL22 and RpL22-like into the testis 80S and ovary 80S cryo-EM maps. The model-to-map fit cross-correlation coefficients are also shown (a higher value corresponds with a better fit). (B) Example of the fit of 2 pairs of equivalent residues for RpL22 into the ovary 80S map, and for RpL22-like into the testis 80S map. (C) Schematic result of the fit of RpL22-like into the testis 80S map, and of RpL22 into the ovary 80S map. For each pair of residues that is different between RpL22 and RpL22-like, the one with the best fit is shown in green. If both residues had a similar fit, then the pair is shown in orange (i.e. the fit was inconclusive).

**Sup 13:**
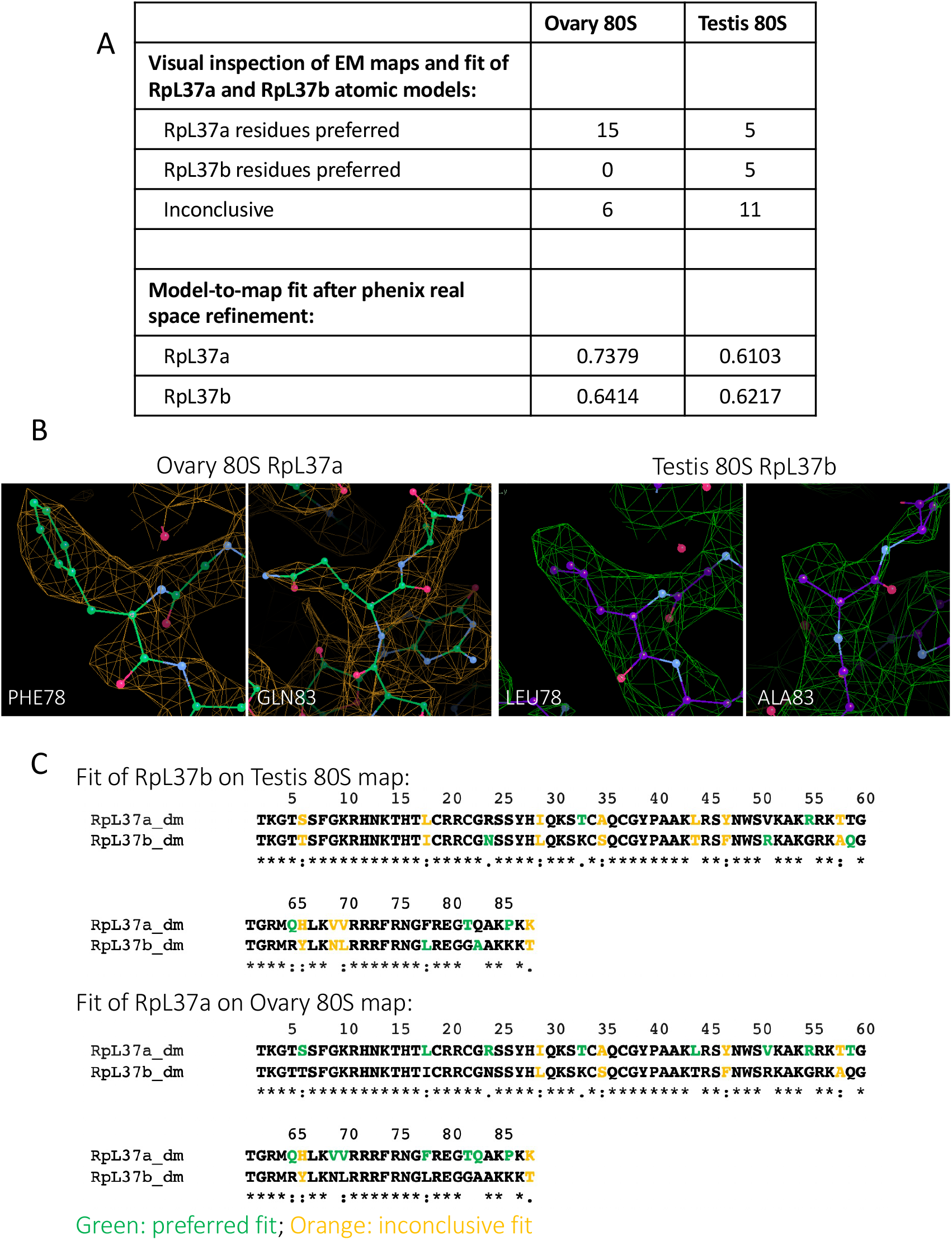
In-depth analysis of density of RpL37a/37b in ovary and testis 80S EM maps. (A) Summary of the fit of residues for RpL37a and RpL37b into the testis 80S and ovary 80S cryo-EM maps. The model-to-map fit cross-correlation coefficients are also shown (a higher value corresponds with a better fit). (B) Example of the fit of 2 pairs of equivalent residues for RpL37a into the ovary 80S map, and for RpL37b into the testis 80S map. (C) Schematic result of the fit of RpL37b into the testis 80S map, and of RpL37a into the ovary 80S map. For each pair of residues that is different between RpL37a and RpL37b, the one with the best fit is shown in green. If both residues had a similar fit, then the pair is shown in orange (i.e. the fit was inconclusive).

**Sup 14:**
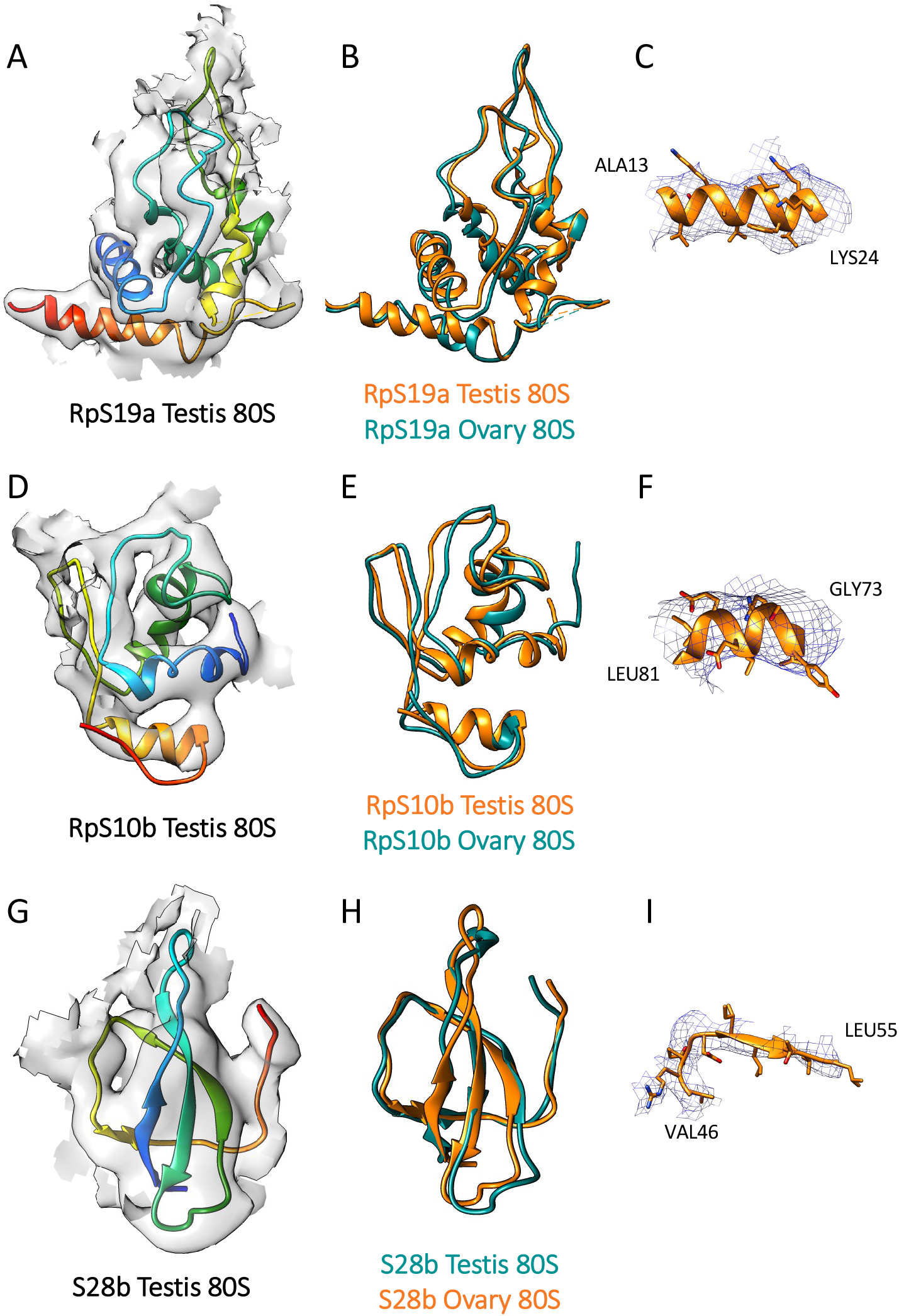
Atomic models of non-switched paralogs with large RMSD between the ovary and testis 80S atomic models. Non-switched paralogs in testis 80S vs ovary 80S are shown. (A-C) RpS19a; (D-F) RpS10b; and (G-I) RpS28b. (A, D and G) Testis atomic models fitted into the EM density. Models are rainbow colored from N-terminus (blue) to C-terminus (red). (B, E and H) Comparison between the testis 80S (orange) and the ovary 80S (teal) atomic models. (C, F and I) Representative fits of the testis 80S atomic models into the EM map.

**Sup 15:**
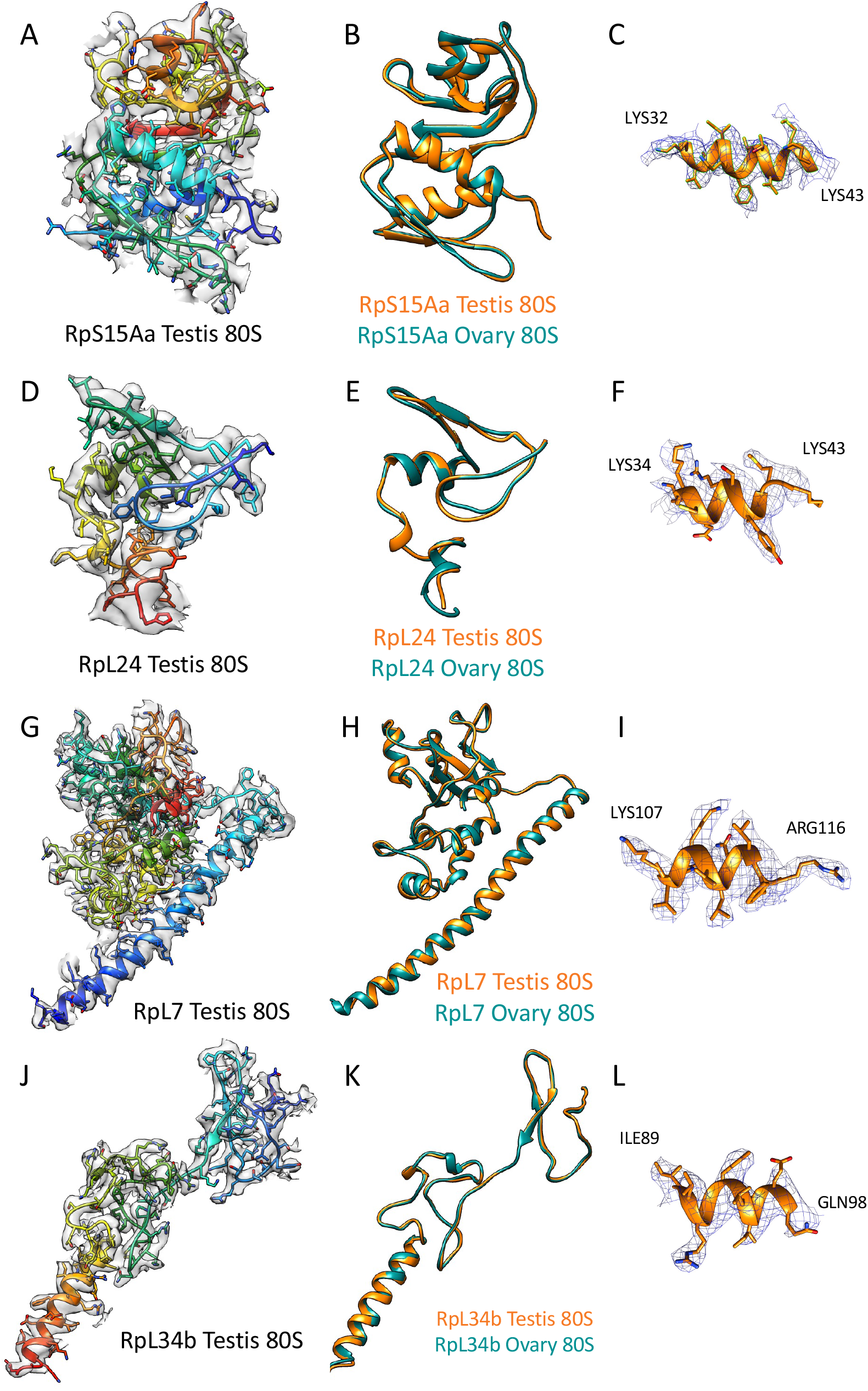
Atomic models of non-switched paralogs with low RMSD between the ovary and testis 80S atomic models. Non-switched paralogs in testis 80S vs ovary 80S are shown. (A-C) RpS15Aa; (D-F) RpL24; (G-I) RpL7; and (J-L) RpL34b. (A, D, G and J) Testis atomic models fitted into the EM density. Models are rainbow colored from N-terminus (blue) to C-terminus (red). (B, E, H and K) Comparison between the testis 80S (orange) and the ovary 80S (teal) atomic models. (C, F, I and L) Representative fits of the testis 80S atomic models into the EM map.

**Sup 16:**
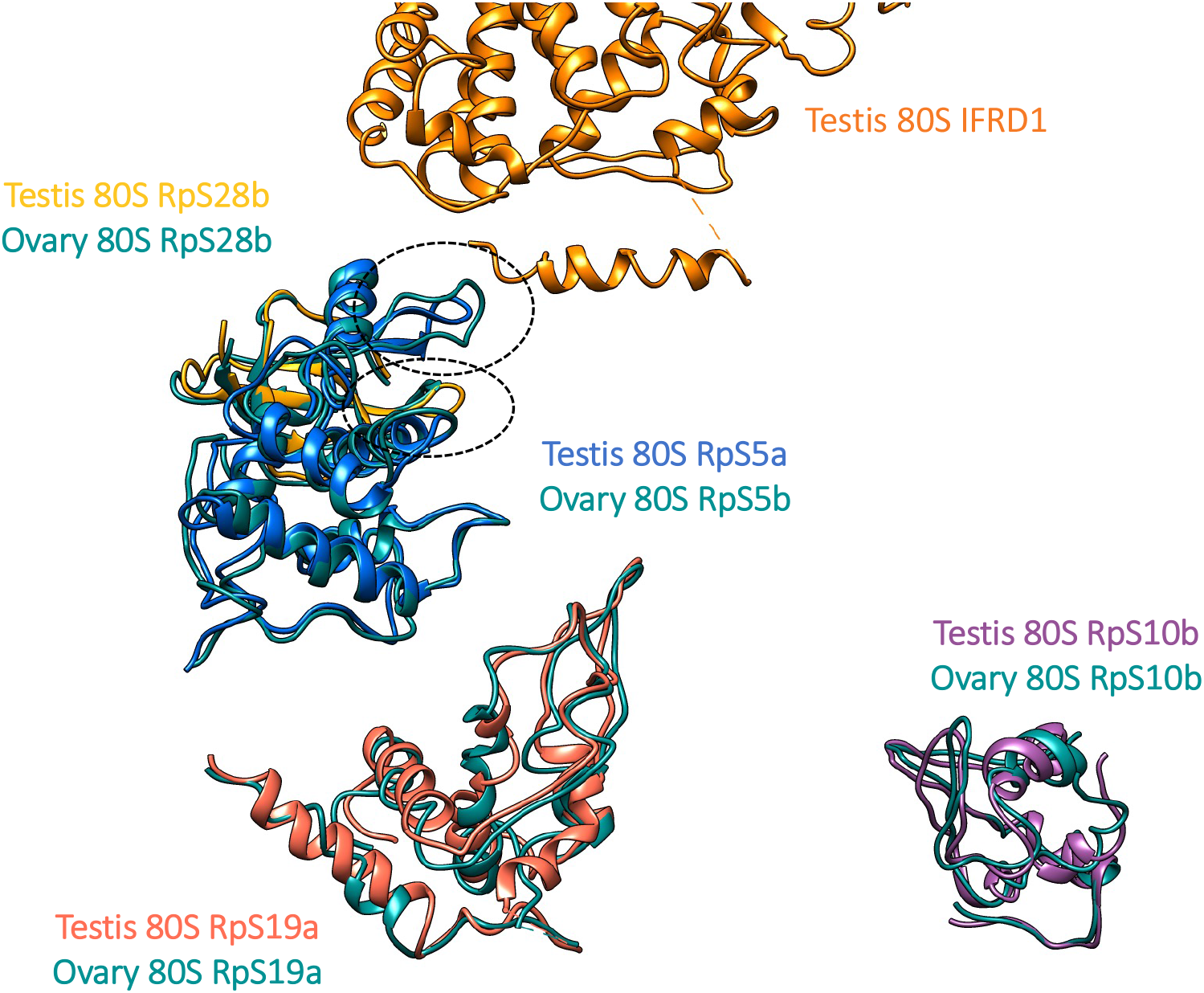
Area around mRNA channel, which in testis 80S is occupied by an alpha-helix from IFRD1. RpS28b (gold), RpS5a (blue) are close to IFRD1 (orange). RpS19a (coral) and RpS10b (purple) are also at the head of the small subunit. Ovary 80S paralogs are superimposed, in teal. The main differences between the PDB models, circled, are in regions close to IFRD1.

**Sup Table 2:**
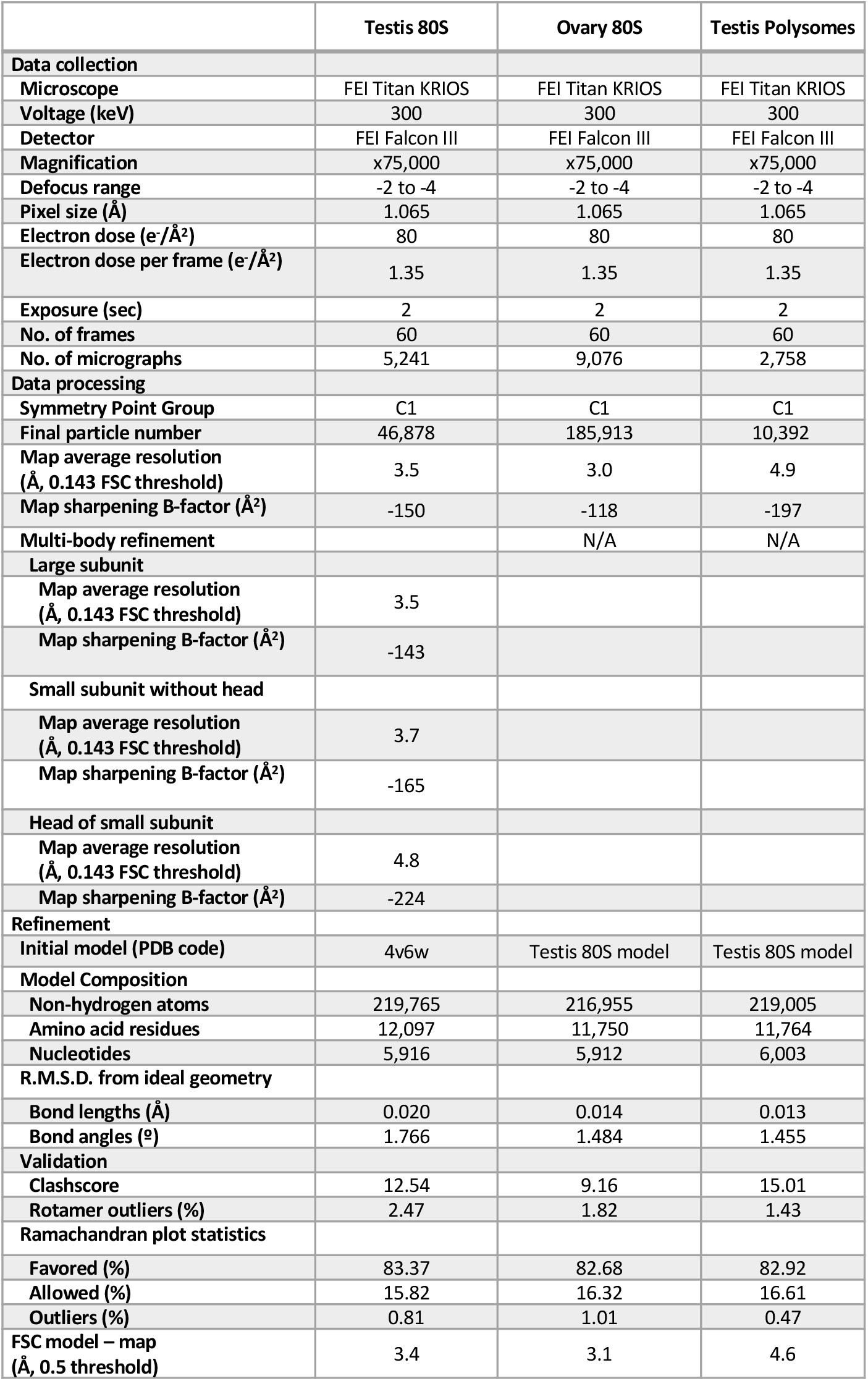
Summary of data collection, image processing, model building, refinement and validation statistics.

